# Functional impairment of “helpless” CD8^+^ memory T cells is transient and driven by prolonged but finite cognate antigen presentation

**DOI:** 10.1101/2024.01.22.576725

**Authors:** Verena van der Heide, Bennett Davenport, Beatrice Cubitt, Vladimir Roudko, Daniel Choo, Etienne Humblin, Kevin Jhun, Krista Angeliadis, Travis Dawson, Glaucia Furtado, Alice Kamphorst, Rafi Ahmed, Juan Carlos de la Torre, Dirk Homann

## Abstract

Generation of functional CD8^+^ T cell memory typically requires engagement of CD4^+^ T cells. However, in certain scenarios, such as acutely-resolving viral infections, effector (T_E_) and subsequent memory (T_M_) CD8^+^ T cell formation appear impervious to a lack of CD4^+^ T cell help during priming. Nonetheless, such “helpless” CD8^+^ T_M_ respond poorly to pathogen rechallenge. At present, the origin and long-term evolution of helpless CD8^+^ T cell memory remain incompletely understood. Here, we demonstrate that helpless CD8^+^ T_E_ differentiation is largely normal but a multiplicity of helpless CD8 T_M_ defects, consistent with impaired memory maturation, emerge as a consequence of prolonged yet finite exposure to cognate antigen. Importantly, these defects resolve over time leading to full restoration of CD8^+^ T_M_ potential and recall capacity. Our findings provide a unified explanation for helpless CD8^+^ T cell memory and emphasize an unexpected CD8^+^ T_M_ plasticity with implications for vaccination strategies and beyond.

## INTRODUCTION

The dynamic regulation of primary (I°) immune cell responses to acute infections or vaccination involves a broad array of tissues, cell types, and extracellular factors that operate in a concerted manner to promote antigen clearance and establishment of immunological memory. CD4^+^ T cells have long been acknowledged to exert a multifaceted role in this process including provision of “help” for expeditious formation of protective CD8^+^ T cell memory^1–6^. While specific requirements may vary, many effector CD8^+^ T cell (T_E_) responses to pathogens, including influenza, lymphocytic choriomeningitis virus (LCMV), vaccinia virus, and *Listeria monocytogenes*, are thought to be helper CD4^+^ T cell-independent^3^, presumably a result of direct antigen-presenting cell activation^7–9^. Yet, despite seemingly normal effector capacity, helpless memory CD8^+^ T cells (T_M_) generated in the absence of CD4^+^ T cells during priming typically expand poorly in response to pathogen rechallenge^10–12^. Accordingly, helpless CD8^+^ T cell memory has been commonly considered a circumscribed scenario in which an integral contribution of CD4^+^ T cell help to initial CD8^+^ T_E_ priming appears to selectively manifest during secondary (II°) CD8^+^ T_E_ responses.

A plethora of reports over the last twenty years have proposed various mechanisms that may cause, prevent, or rescue helpless CD8^+^ T cell memory, including contributions by different immune cell populations (*e.g.*, CD4^+^ T_E_, regulatory CD4^+^ T cells, licensed dendritic cells, antibody-producing B cells)^2,4,9,13,14^, co-stimulatory and inhibitory interactions (*e.g.*, via CD40-CD40L or Programmed Cell Death Protein 1/PD-1-PD-L1)^2,4,6,9,14^, transcription factors (*e.g.*, T-bet)^15,16^, cytokines (*e.g.*, IL-2, type I and type II interferons)^2,4,13,14,16^ as well as alterations of T cell migration^4,17^, metabolism^18^, and epigenetic properties^19,20^. In aggregate, they underscore differential requirements of CD4^+^ T cell-mediated activities across the spectrum of specific CD8^+^ T cell immunity and the notion that helpless priming may promote CD8^+^ T_M_ dysfunction^21–23^. Yet, our understanding of helpless CD8^+^ T cell memory remains incomplete, in part owing to the complexity of interactions between various immune cell types as well as differences across experimental systems studied to date, some of which may in fact not be strictly CD4^+^ T cell-independent, thereby complicating the interpretation of at times seemingly incompatible reported results.

Here, we revisit the phenomenon of helpless CD8^+^ T cell memory in the context of its extended temporal organization. Phenotypically and functionally heterogeneous, long-lived CD8^+^ T_M_ are not a fixed entity but continue to evolve over time both at the population and the subset level, including a compositional shift from terminally differentiated cytotoxic “effector memory” CD8^+^ T_EM_ to quiescent pluripotent “central memory” T_CM_^24–31^. This developmental path involves profound transcriptional, phenotypic, and functional remodeling pertaining to expression levels of markers associated with memory formation (*e.g.*, CXCR3, CX3CR1, KLRG1, CD127)^26,28,29,32–34^, redistribution of CD8^+^ T_M_ to secondary lymphoid organs (SLOs)^29,34^, metabolic adaptations^29,35^ as well as generally improved functionality and long-term survival^27–29^. Notably, only a subset of these pathways are mechanistically linked to improved CD8^+^ T_M_ responsiveness^30^. However, in conjunction they correlate with an enhanced II° CD8 T_E_ proliferative potential as well as more efficient immune protection against rechallenge^24,27,28,30^ and, similar to observations for yellow fever vaccine-specific human CD8^+^ T_M_^36–38^, they align with a broad “dedifferentiation” program imparting gradual acquisition of a more homogenous “naive-like” CD8^+^ T_M_ phenotype while reinforcing a “mature” polyfunctional CD8^+^ T_M_ core signature^28,39^.

Accordingly, we hypothesized that the helpless CD8^+^ T_M_ recall deficit might not be a permanent defect but rather reflect a transient delay of CD8^+^ T_M_ maturation. Working with the LCMV model of acute viral infection, we find that CD8^+^ T_E_ priming in the absence of CD4^+^ T cells allows for broadly normal transcriptional, phenotypic, and functional CD8^+^ T_E_ differentiation including effective generation of memory precursor (MP) CD8^+^ T_E_. We identify prolonged but finite exposure of developing and established helpless CD8^+^ T_M_ to cognate antigen as a proximate cause for helpless CD8^+^ T cell memory and, importantly, demonstrate that the associated phenotypic and functional defects effectively resolve over time resulting in complete restoration of CD8^+^ T_M_ recall potential. Our observations thus redefine helpless CD8^+^ T cell memory as a temporary rather than a terminal state with implications for our general understanding of CD8^+^ T cell immunity specifically with respect to vaccination protocols but also in autoimmunity and cancer where CD8^+^ T cell function may be decisively shaped by protracted antigen exposure.

## RESULTS

### Modeling helpless CD8^+^ T cell memory in acutely-resolving LCMV infection

Acute infection of its natural murine host with the LCMV Armstrong (Arm) strain is typically resolved within 8-10 days post infection (dpi) in a CD8^+^ T cell-dependent manner that does not appear to require contributions by CD4^+^ T cells^40–44^. To revisit helpless CD8^+^ T cell memory in this host-pathogen system, we generated T cell receptor (TCR)-transgenic P14 chimeras in which a small trace population of naïve congenic LCMV GP_33-41_-specific P14 CD8^+^ T cells (P14 T_N_) is transferred into isotype-treated helped or transiently CD4^+^ T cell-depleted helpless C57BL/6 (B6) recipients before LCMV Arm infection (***Figures 1A and S1A-S1B***). In this setup, P14 T_E_ expansion and contraction kinetics are comparable to endogenously induced LCMV-specific CD8^+^ T cell responses^28,45^, and, importantly, similar between helped and helpless conditions allowing for the establishment of commensurate P14 T_M_ pools ∼7-9 weeks after I° infection (***Figure S1C***). To assess II° P14 T_E_ reactivity, we adoptively transferred enriched helped or helpless P14 T_M_ populations into naïve congenic B6 hosts that were subsequently challenged with high-dose recombinant *Listeria monocytogenes* expressing the LCMV glycoprotein gp_33-41_ determinant (rLM-gp33) or with the original LCMV Arm strain (***Figure 1A***). As anticipated, in comparison with their helped counterparts, helpless P14 T_M_ mounted a significantly smaller recall response to either infectious agent in peripheral blood, spleen, and lymph nodes (LNs) (***Figures 1B and S1D***), indicating that the impaired II° effector expansion by helpless P14 T_M_ was agnostic to the type of rechallenge pathogen. Thus, our model recapitulates the cardinal feature of helpless CD8^+^ T cell memory^10–12^ and further emphasizes that the resulting defect is likely the consequence of altered properties intrinsic to helpless P14 T_M_.

**Figure 1.**
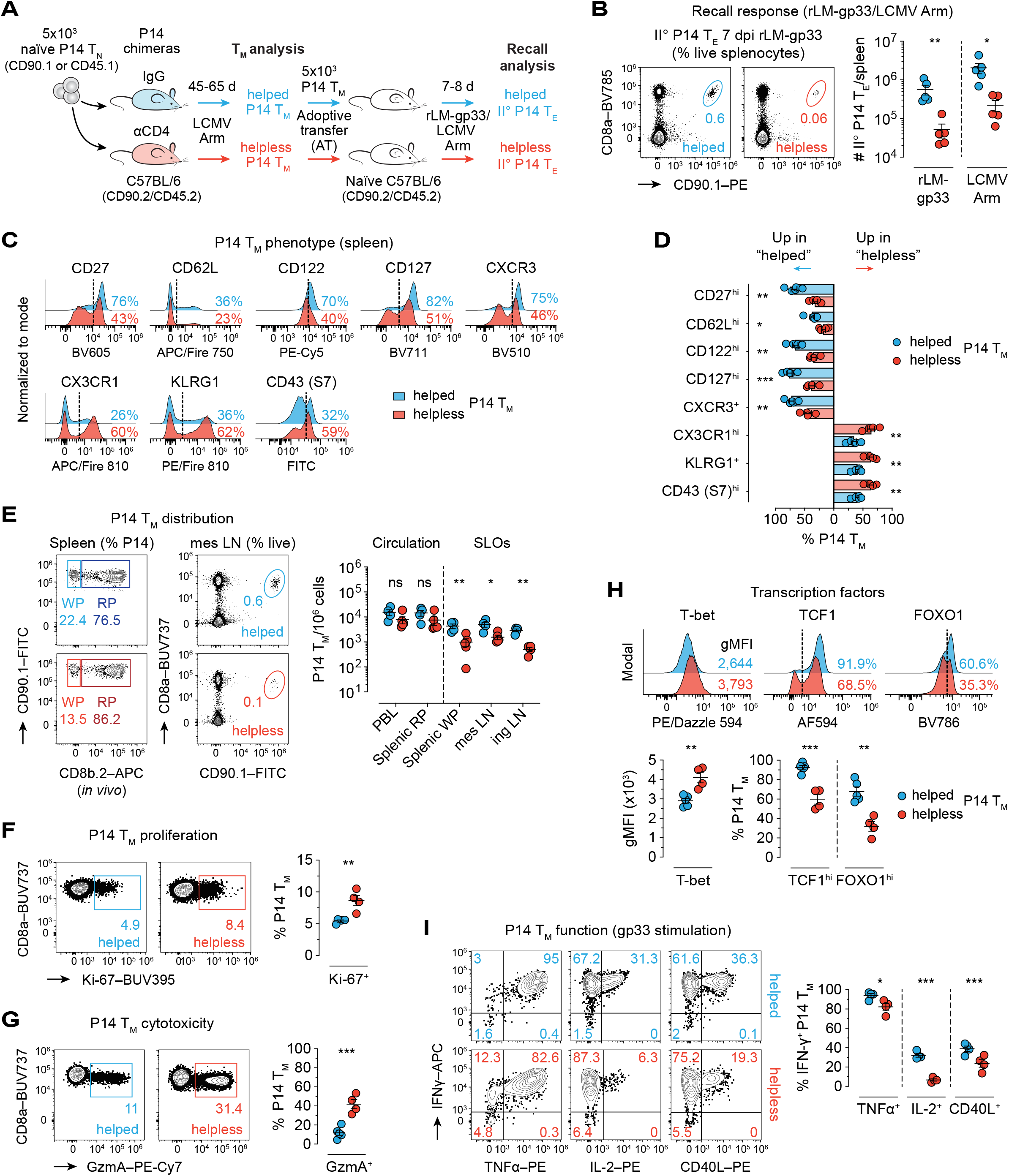
Phenotypical and functional changes in helpless CD8^+^ T_M_ indicate broadly impaired memory maturation. (**A**-**I**) Comparative analysis of early helped and helpless CD8^+^ T_M_ (**A**) LCMV-immune helped or helpless P14 chimeras were generated by transferring naïve congenic P14 (P14 T_N_) cells into isotype (IgG)-treated or transiently CD4^+^ T cell-depleted (αCD4) naïve hosts followed by acute LCMV Arm infection. Memory P14 cells (P14 T_M_) were phenotypically and functionally evaluated ∼45-65 days (d) later. (**B**) (Left) representative flow cytometry plots of P14 recall response to rLM-gp33 and (right) enumeration of splenic secondary P14 effector (II° P14 T_E_) expansion after rLM-gp33 or LCMV Arm rechallenge (see ***Figure S1D***). (**C**-**D**) (**C**) Representative histogram overlays of distinctive helped and helpless spleen P14 T_M_ phenotypes and (**D**) corresponding statistical summary. (**E**) (Left) representative flow cytometry plots of P14 T_M_ distribution across splenic white (WP) and red pulp (RP) as well as mesenteric lymph nodes (mes LNs) and (right) P14 numbers in circulation (peripheral blood/PBL, RP) and selected secondary lymphoid organs (SLOs: WP, mes and inguinal/ing LNs). (**F**-**H**) Representative flow cytometry plots/histogram overlays and statistical analyses of splenic P14 T_M_ (**F**) proliferation, (**G**) cytotoxicity as well as (**H**) T cell-differentiation-associated transcription factor profiles. (**I**) Representative flow cytometry plots and quantification of P14 T_M_ after *in vitro* gp33 peptide stimulation. Data representative of one out of two-three independent experiments with n=4-5 mice/group. Graphs show individual helped (blue) and helpless (red) mice with mean±SEM. \**p*<0.05, \*\**p*<0.01, and \*\*\**p*<0.001 by unpaired Student’s *t* test or Mann Whitney U test. ns, non-significant; AT, adoptive transfer; dpi, days post infection.

### Helpless CD8^+^ memory T cells are broadly immature

Since CD8^+^ T_M_ recall capacity correlates with a progressive acquisition of mature T_M_ properties^24,25,27,28,30^, we hypothesized that the compromised II° effector response of helpless P14 T_M_ might be associated with phenotypic and functional alterations consistent with an impaired maturation process. In alignment with our hypothesis, we observed that helpless splenic P14 T_M_ evaluated ∼7-9 weeks after acute LCMV infection exhibited an immature phenotype with increased expression of “terminal differentiation”^32–34,46–49^ markers, including CX3CR1, KLRG1, KLRC1, CD43, and CD54, while helped P14 T_M_ featured higher levels of traditional “memory” molecules^32,34,39,48–51^, such as CD27, CD62L, CD127, CXCR3, CCR7, Ly6C, Ly6A/E, and Slamf6 (***Figures 1C-1D and S1E***).

In recent years, the originally proposed classification^52^ into lymphoid tissue-bound CD62L^hi^CCR7^+^ T_CM_ and systemically trafficking CD62L^lo^CCR7^-^ T_EM_ has been expanded by a variety of markers to capture existing granularity among circulating CD8^+^ T_M_ and to better distinguish broadly “multipotent” (*e.g.*, CD62L^hi^CD27^hi^CD127^hi^CD43^lo^KLRG1^lo^ or CD62L^+^CD27^+^CXCR3^+^CX3CR1^-^ CD8^+^ T_CM_) from functionally heterogenous “transitional effector-like” (including CD62L^-^ CXCR3^+^CD127^+^CX3CR1^-^, CD62L^-^CXCR3^-^CD127^+^CX3CR1^+^, and KLRG1^hi^CD127^int^CD27^lo^ CD8^+^ T_EM_) as well as “terminally differentiated” subsets (*e.g.*, CD62L^-^CD127^-^CXCR3^-^CX3CR1^+^ CD8^+^ T_t-EM_)^32,46,48,49,53–55^. Principal component analysis (PCA) guided by these markers segregated helped and helpless P14 T_M_ into two clusters, underlining phenotypic diversity between both groups (***Figure S1F***). This was further mirrored by a relative increase in transitional effector-like and terminally differentiated P14 T_EM_ yet fewer T_CM_ in peripheral blood and the splenic red pulp (***Figure S1G***). In contrast, the subset composition of helped and helpless P14 T_M_ in the splenic white pulp or mesenteric LNs was indistinguishable (***Figure S1G***), even though total helpless P14 T_M_ numbers in these compartments were reduced (***Figure 1E***) likely as a result of delayed SLO homing/tropism^29,34^. Simultaneously, however, bulk frequencies of helped and helpless circulating P14 T_M_ in peripheral blood and the splenic red pulp remained remarkably similar (***Figure 1E***).

We further detected increased Ki-67 and granzyme A (GzmA) levels in helpless P14 T_M_ pointing to a more recent history of proliferation and cytotoxic activity (***Figures 1F-1G***). Similarly, higher T-bet – as reported previously^15^ – yet reduced TCF1 and FOXO1 expression further corroborated an impaired T_M_ maturation process, consistent with the respective roles of these transcription factors in CD8^+^ T_E_^56^ and CD8^+^ T_M_ generation^57,58^ (***Figure 1H***). Limited helpless P14 T_M_ polyfunctionality after *in vitro* gp33-peptide restimulation equally conformed to an earlier CD8^+^ T_M_ stage (***Figure 1I***), and led us to hypothesize that acquisition of a mature phenotype by helpless P14 T_M_ as well as associated functions may occur at a delayed pace. Indeed, phenotypic clustering of helped and helpless P14 T_M_ in peripheral blood and SLOs along with differential TCF1 expression provided preliminary evidence for the potential temporal nature of helpless CD8^+^ T cell memory, indicating an expedient capacity for helpless T_M_ maturation in restricted anatomic compartments whereas helpless features largely pertained to circulating P14 T_M_ (***Figures S1H-S1I***).

Intriguingly, since homeostatic proliferation of CD8^+^ T_M_ remains stable over time^29^, increased Ki-67 expression in conjunction with enhanced GzmA by helpless P14 T_M_ (***Figures 1F-1G***) suggested a potential antigen-driven component although the helpless phenotype in our model is fundamentally distinct from the presentation of *bona fide* exhausted virus-specific CD8^+^ T cells (T_EX_) in persistent LCMV clone 13 (cl13) infection^59^ (***Figure S1J***).

### Helpless CD8^+^ T_M_ cells undergo delayed phenotypic maturation

To evaluate the potential temporality of the observed helpless CD8^+^ T_M_ defects, we monitored relative P14 T_M_ numbers in peripheral blood of helped *vs*. helpless P14 chimeric cohorts for >1 year and assessed the longitudinal evolution of markers originally distinguishing helped and helpless P14 T_M_ at the early memory stage (***Figures 2A-2B***). As expected^28,34,48^, helped P14 T_M_ gradually upregulated CD27, CD62L, CD122, CD127, and CXCR3, while expression of CX3CR1, KLRG1, and CD43 (115 kDa glycoform) was concurrently downmodulated (***Figure 2C***). Interestingly, helpless P14 T_M_ were maintained at comparable frequencies and displayed similar maturation trajectories, albeit with considerably delayed kinetics acquiring a “helped-like” mature T_M_ phenotype on average only >1 year after LCMV infection (**Figures 2B-2C**). As a result, phenotypic diversity between helped and helpless P14 T_M_ peaked at ∼120 dpi for most of the above markers except for CD127 (∼100 dpi) and CD122 (∼220 dpi) (***Figure S2A***). Yet, despite distinct phenotypical compositions four months after LCMV (***Figure S2B***), the associated difference in recall capacity, while still significant, was comparatively reduced, indicating that phenotypic evolution of helpless P14 T_M_ trails behind a potential equilibration of II° P14 T_E_ responses *in vivo* (***Figure S2C and see Figure 1B***). Additionally, the relatively homogenous maturation of helped P14 T_M_ contrasted with greater variability among helpless P14 T_M_ suggesting that P14 T_M_ kinetics are shaped by processes that operate with differential efficacy in individual helpless mice.

**Figure 2.**
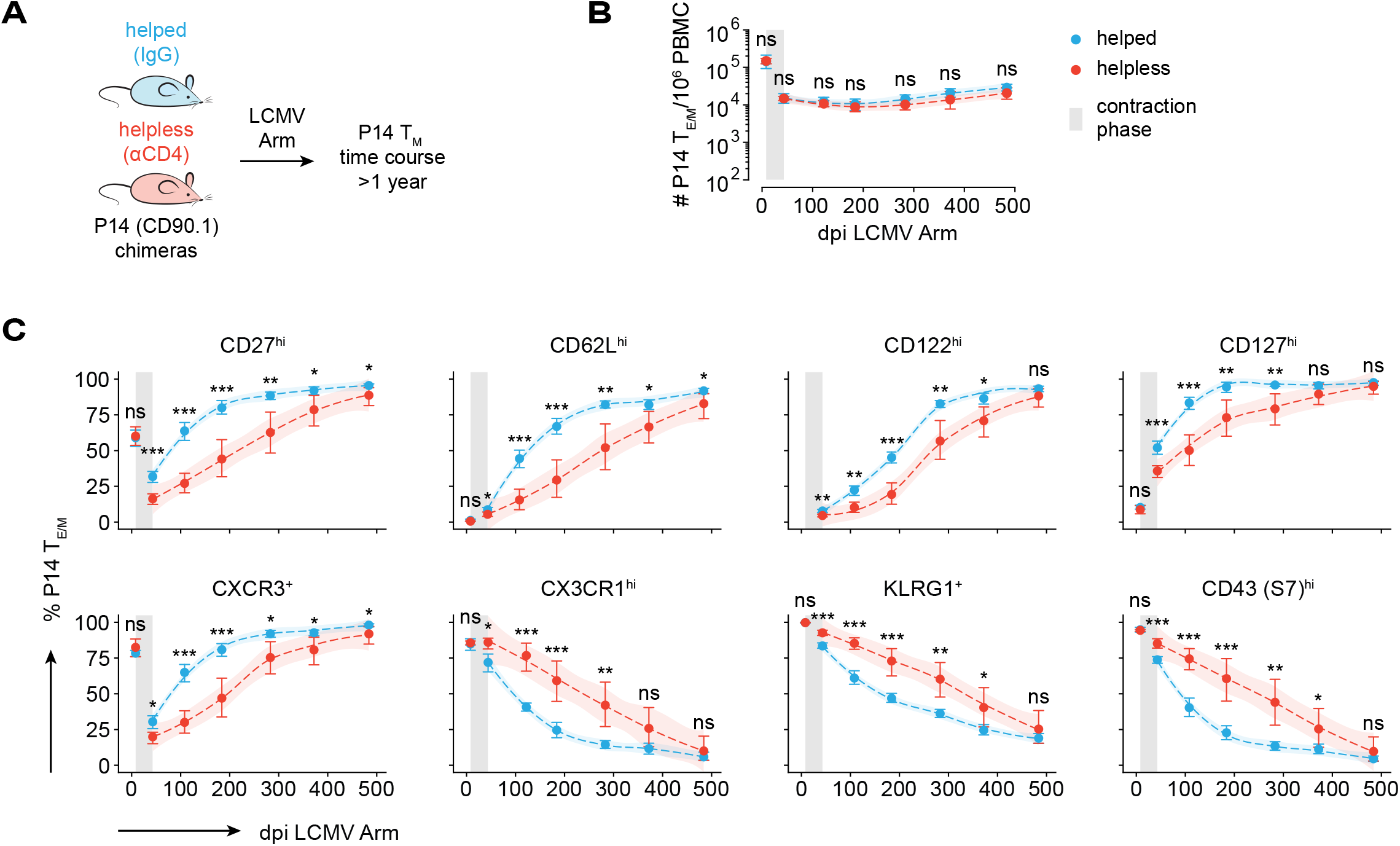
Helpless CD8^+^ T_M_ are subject to delayed phenotypic memory maturation. (**A**-**C**) Time course analysis of helped and helpless P14 T_M_ among peripheral blood mononuclear cells (PBMC). (**A**) Experimental setup in the transient CD4^+^ T cell depletion model. (**B**-**C**) (**B**) Maintenance of P14 T_M_ and (**C**) progressive phenotypic maturation from peak of effector phase (8 dpi) throughout long-term memory. Filled symbols and error bars indicate mean±bootstrap 95% confidence intervals. Connecting dashed lines and ribbons represent LOESS smoothing trends with 95% confidence intervals calculated by the geom_smooth() function in R. n=5-14 mice/group/time point. Data for d8 P14 T_E_ collected from a separate cohort of mice and added for visual context. \**p*<0.05, \*\**p*<0.01, and \*\*\**p*<0.001 by unpaired Student’s *t* or Mann Whitney U test. ns, non-significant.

### Helpless CD8^+^ T cell memory defects are resolved in aged CD8^+^ T_M_

The preceding results suggested that the eventual acquisition of selected mature memory traits by helpless P14 T_M_ may be accompanied by a broader resolution of the multiple helpless CD8^+^ T_M_ defects recorded in the early memory phase (***Figures 1 and S1***). Mirroring our observations in peripheral blood, we found that despite some variability, splenic P14 T_M_ were maintained at proportionate numbers in helped and helpless P14 chimeras at >450 dpi (***Figure 3A***). Strikingly, helped and helpless P14 T_M_ now presented with near-identical expression patterns of molecules that distinguished both groups at earlier time points (***Figures 3B-3C and S2D***), hence aligning aged helped and helpless P14 T_M_ clusters and demarcating them against their early-memory-stage equivalents (***Figures 3D and S2E***). Similarly, we noted an identical distribution of helped and helpless P14 T_M_ across various SLOs, including the splenic white pulp, which was accompanied by equivalent P14 T_M_ numbers in these tissues in both groups (***Figures 3E and S2F***).

**Figure 3.**
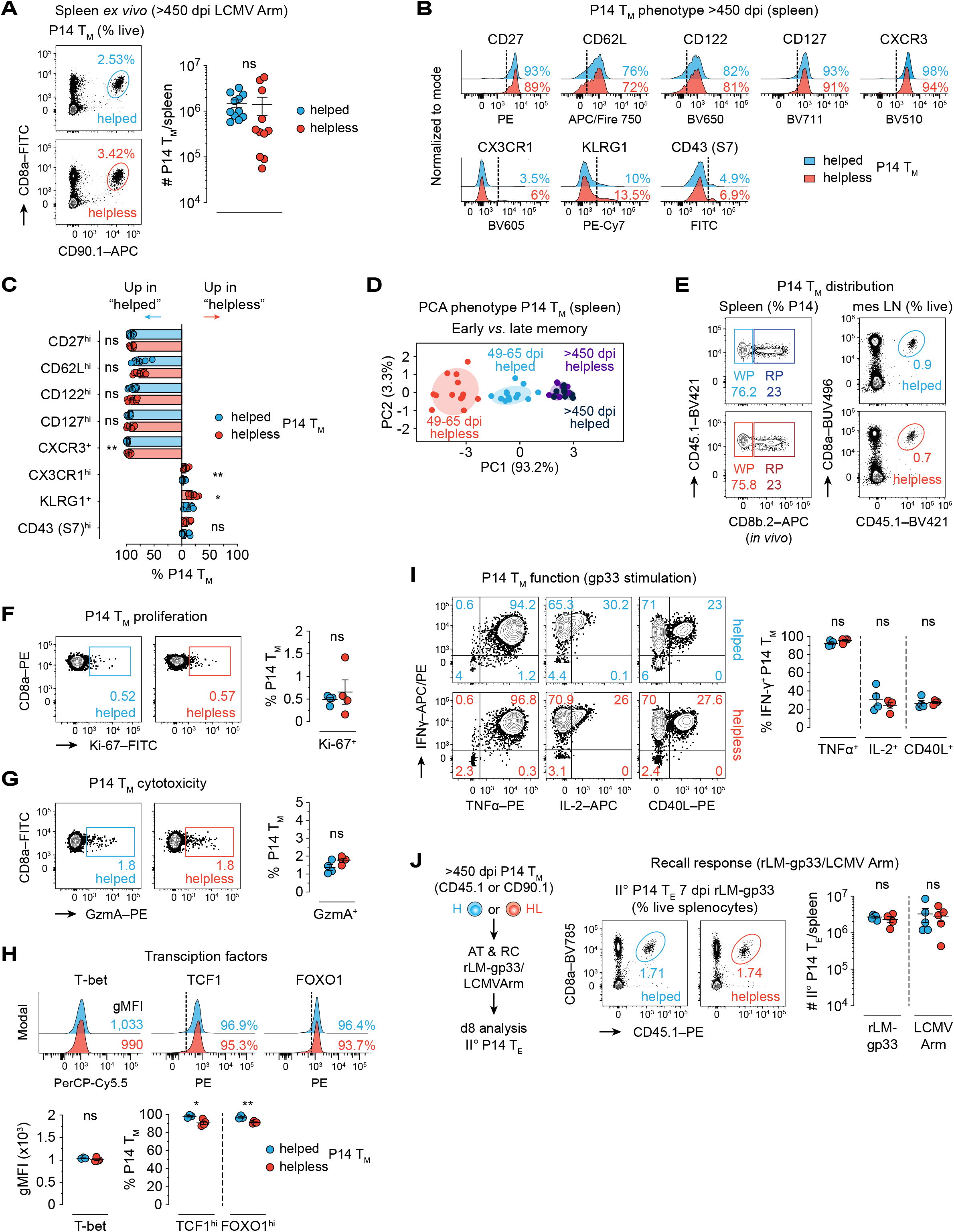
Helpless memory defect is comprehensively resolved in aged CD8^+^ T_M_. (**A**-**J**) Comparative analysis of helped and helpless CD8^+^ T_M_ >450 dpi (transient CD4^+^ T cell depletion model) (see Figure 1). (**A**) (Left) representative flow cytometry plots and (right) splenic P14 T_M_ enumeration. (**B**-**C**) (**B**) Representative histogram overlays of helped and helpless splenic P14 T_M_ phenotypes and (**C**) corresponding statistical summary. (**D**) PCA based on percentual expression of markers in (**B**; with exception of CD122) comparing early (49-65 dpi) *vs*. late (>450 dpi) helped and helpless P14 T_M_ (see ***Figures S1F*** and ***S2E***). (**E**) P14 T_M_ distribution across splenic WP, RP, and mes LNs by representative flow cytometry plots (see ***Figure S2F***). (**F**-**H**) Representative histogram overlays/flow cytometry plots of (**F**) splenic P14 T_M_ proliferation, (**G**) cytotoxicity, and (**H**) transcription factor profiles. (**I**) P14 T_M_ functionalities. (**J**) (Left) representative flow cytometry plots and (right) quantification of aged splenic II° P14 T_E_ response to rLM-gp33/LCMV Arm rechallenge (RC) (see ***Figure S2G***). Data pooled from three independent experiments in (**A**)-(**C**) and representative of one out of two independent experiments in (**E**)-(**J**) with n=3-5 mice/group/experiment. PCA with 80% confidence ellipses in (**D**) combined from two independent experiments conducted at 49-65 dpi and three independent experiments at >450 dpi. Graphs show individual helped (blue) and helpless (red) mice with mean±SEM. \**p*<0.05, \*\**p*<0.01 by unpaired Student’s *t* test or Mann Whitney U test. ns, non-significant.

Commensurate Ki-67 and loss of *ex vivo* detectable GzmA expression by helped and helpless P14 T_M_ further indicated matching homeostatic properties (***Figures 3F-3G***) and together with comparable T-bet content pointed to a now overall aligned T_M_ maturation stage (***Figure 3H***). This conclusion was also supported by enhanced levels of the memory-associated transcription factors TCF1 and FOXO1 in helpless P14 T_M_, lagging only slightly behind their helped counterparts (***Figure 3H***). In addition, diversified functional profiles of helpless P14 T_M_ after gp33-peptide restimulation now matched those of helped P14 T_M_ and suggested complete functional maturation of helpless P14 T_M_ (***Figure 3I***). Most importantly, we found that the systemic II° expansion of helped and helpless P14 T_M_ in response to either rLM-gp33 or LCMV Arm rechallenge was now enhanced and indistinguishable (***Figures 3J and S2G***), thus confirming our hypothesis that helpless CD8^+^ T cell memory is indeed a temporary rather than terminal defect that may effectively resolve over time.

To extend these observations beyond the TCR-transgenic P14 chimera system, we interrogated the phenotypic maturation of endogenously generated LCMV-specific D^b^GP_33_^+^ CD8^+^ T_M_ in peripheral blood of IgG-treated *vs*. transiently CD4^+^ T cell-depleted LCMV-immune B6 mice. Consistent with our P14 data, pronounced differences between helped and helpless virus-specific CD8^+^ T_M_ still present at ∼200 dpi mostly resolved by ∼2 years after acute LCMV (***Figure S2H***). Notably, notwithstanding differential T_M_ maturation dynamics between both groups, helped and helpless D^b^GP_33_^+^ CD8^+^ T_M_ appeared to trail behind their P14 counterparts by several months, presumably due to lower frequencies of LCMV-specific precursor T_N_ resulting in an overall associated delay of CD8^+^ T_M_ maturation kinetics^28,45,60,61^.

### Helped and helpless CD8^+^ T_E_ cells share a common transcriptomic signature

A proximate cause for the development of helpless memory may be altered CD8^+^ T_E_ differentiation. Yet, earlier reports^10–12^ and our data on bloodborne P14 T_E_ (***Figures 2B-2C***) did not reveal any gross numerical or phenotypic differences between helped and helpless P14 chimeric mice at the peak of the effector response on day 8 after LCMV Arm infection. Furthermore, P14 T_E_ expansion kinetics in the spleen as well as associated subset distribution into CD127^-^KLRG1^-^ early (EEC), CD127^-^KLRG1^+^ short-lived (SLEC), CD127^+^KLRG1^-^ memory precursor (MPEC), and CD127^+^KLRG1^+^ double-positive effector cells (DPEC), respectively^56,62–64^, were indistinguishable between both groups with the notable exception of a trending increase in helpless P14 EECs at 5 dpi as well as a subtle MPEC reduction at 5 and 6.5 dpi, suggesting slightly delayed T_E_ differentiation (***Figures 4A-4C***). In apparent contrast, published transcriptomic profiling of helped *vs.* helpless LCMV-specific CD8^+^ T_E_ by RNA sequencing (RNA-seq) demonstrated distinct differences even though corresponding protein expression studies were not performed^65^.

**Figure 4.**
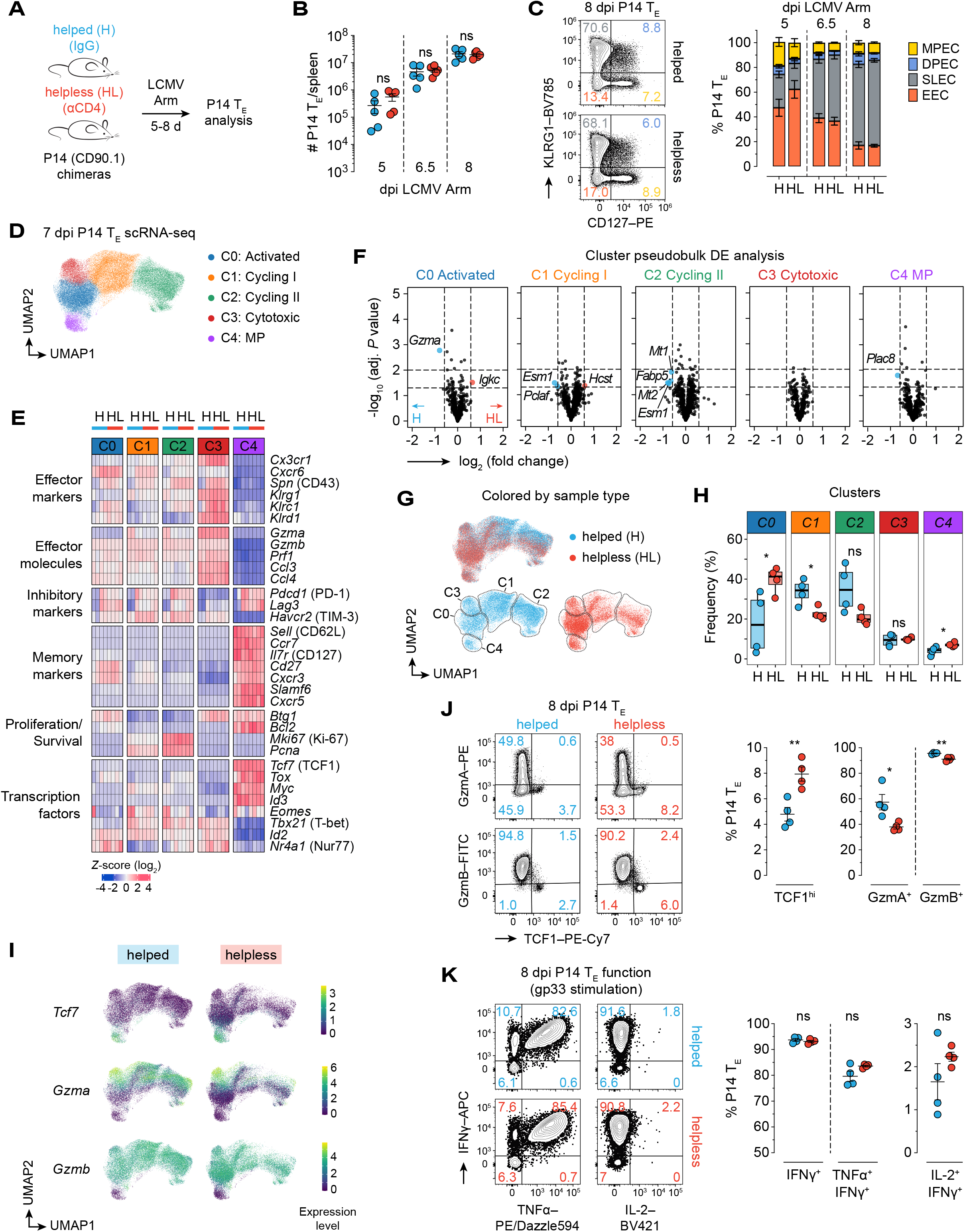
Helpless CD8^+^ T_E_ differentiation is only minimally altered. (**A**-**K**) Analysis of helped (H) and helpless (HL) splenic P14 T_E_ differentiation. (**A**) Experimental design in the transient CD4^+^ T cell depletion model. (**B**-**C**) (**B**) Splenic P14 T_E_ quantification 5-8 dpi and (**C**) flow-cytometric evaluation of early effector (EEC), short-lived effector (SLEC), double-positive effector (DPEC), and memory precursor effector (MPEC) subsets based on KLRG1 and CD127 expression (left) and summary analysis (right). (**D-I**) scRNA-seq analyses of 7 dpi helped and helpless P14 T_E_ (see ***Figure S3*** and ***Table S1***). (**D**) UMAP projection with five Leiden clusters. (**E**) Heatmap of selected *Z*-score log_2_-normalized marker gene expression across clusters stratified by helped and helpless P14 T_E_ status. (**F**) Differential expression (DE) cluster pseudobulk analysis between splenic helped and helpless P14 T_E_. Vertical dashed lines indicate 1.5-fold change; horizontal dashed lines mark *p*=0.05 and 0.01, respectively. (**G**-**H**) Relative differential cluster abundance (**G**) visualized by UMAP plots colored by sample type and (**H**) statistical evaluation between helped and helpless conditions. (**I**) Feature plots of *Tcf7*, *Gzma*, and *Gzmb* mRNA expression across clusters and helped *vs*. helpless experimental conditions. (**J**) (Left) representative flow cytometry plots of TCF1^+^ memory precursor (MP) *vs*. GzmA^+^ or GzmB^+^ effector-phenotype 8 dpi splenic P14 T_E_ and (right) statistical analysis. (**K**) Flow-cytometric analysis of P14 T_E_ function after gp33-peptide restimulation (left) and (right) statistical assessment. Helped (n=4) and helpless P14 T_E_ (n=4) in the scRNAseq experiment were sort-purified and processed at the same time as independent samples. Graphs representative of one out of at least two independent experiments plot individual helped (blue) and helpless (red) mice with n=4-5 animals/group/time point/experiment and mean±SEM. Boxplots with individual mice in (**F**) indicate median, quartiles, and range. \**p*<0.05, \*\**p*<0.01 by unpaired Student’s *t* test or Mann Whitney U test. ns, non-significant.

To address this discrepancy, and to account for the fact that bulk RNA-seq cannot readily capture subset skewing, we conducted single-cell RNA-sequencing (scRNA-seq) analyses of helped and helpless P14 T_E_ at 7 dpi as detailed in ***Methods***. Embedding of individual helped and helpless P14 cells into uniform manifold approximation (UMAP^66^) followed by unsupervised Leiden clustering^67^ identified five subpopulations that were annotated based on respective marker genes as “activated”, “cycling I”, “cycling II”, “cytotoxic” and “memory precursor (MP)”, respectively (***Figures 4D and S3A-S3C; Table S1***). Of note, similar results were obtained by label transfer using a recently published scRNA-seq data set of P14 T_E_, T_M_, and T_EX_ generated in acute LCMV Arm or chronic LCMV cl13 infection^68^ (***Figures S3D-S3F***). Interestingly, we found that mRNA profiles of multiple genes typically linked to CD8^+^ T_E_ differentiation and/or CD8^+^ T_M_ generation, including *Cx3cr1, Gzma* (encoding granzyme A)*, Gzmb* (encoding granzyme B)*, Sell* (encoding CD62L), and *Tcf7* (encoding TCF1), were indistinguishable between the respective helped and helpless clusters (***Figure 4E***). This observation was further corroborated in complementary cluster pseudobulk analyses^69,70^ demonstrating a near-complete absence of differentially expressed genes (DEGs) between helped and helpless P14 T_E_ with the notable exception of higher *Gzma* levels in “activated” helped P14 T_E_ (***Figure 4F***). Similarly, combining all clusters for global pseudobulk analyses revealed only minor differences (*i.e.,* differential expression of 12/526 genes) collectively pointing towards a slight enhancement of cell cycle-associated gene expression (*Knl1, Esco2, Clspn, Rrm2, Pcla*) by helped P14 T_E_ (***Figures S3G-S3H***).

Instead, distinctive differences pertained to differential cluster abundance including a relative increase of helpless P14 T_E_ in the “activated” cluster, fewer “cycling” helpless P14 T_E_, and, unexpectedly, a greater fraction of helpless P14 T_E_ in the “MP” cluster; no differences were recorded for the “cytotoxic” cluster (***Figures 4G-4H and S3I***). We also noted some variability in the size of the “activated” and “cycling” clusters across biological replicates (particularly for helped P14 T_E_) (***Figure S3I***) and, given their considerable similarity (***Figure S3C***), we therefore repeated lower resolution Leiden clustering followed by adjusted UMAP projection. Indeed, this approach collapsed all activated and cycling subpopulations into a single “activated/effector” cluster of comparable size for helped and helpless P14 T_E_, now leaving the greater preponderance of helpless P14 T_E_ MPs as the only significant distinction at the level of differential cluster abundance (***Figures S3J-S3L***).

### Subtle phenotypic skewing of helpless CD8^+^ T_E_ differentiation in favor of MP T_E_ generation

To further leverage distinctive transcriptomic signatures for phenotypic demarcation of P14 T_E_ subsets akin to our lower-resolution UMAP projection (***Figure S3J***), we considered marker genes (***Figure S3C and Table S1***), their expression profiles across clusters (***Figure 4E***), and the suitability to visualize corresponding proteins at the single-cell level. Here, *Tcf7* was largely confined to the “MP” subset while *Gzma* was highly expressed in the “cytotoxic” cluster, yet also found in a substantial fraction of cycling P14 T_E_. In contrast, *Gzmb* was uniformly detected with exception of the “MP” population (***Figure 4I***). Flow-cytometric analysis of helped and helpless P14 T_E_ broadly confirmed these patterns at the protein level – including reciprocal TCF1 *vs*. Granzyme B (GzmB) distribution – revealing a modest yet significant reduction of GzmA^+^ and GzmB^+^ effectors as well as a higher fraction of TCF1^hi^ MP T_E_ in helpless P14 chimeric mice (***Figure 4J***), consistent with a pivotal role of TCF1 for MP formation at the expense of T_E_ differentiation^71–73^. Additional stratification of TCF1^hi^ P14 T_E_ further illustrated that their preferential albeit not exclusive MPEC phenotype accounted for the greater MP abundance among helpless P14 T_E_ (***Figure S4A***). Notably, we also observed a trend towards higher frequencies of helpless CD62L^hi^ MPECs (***Figure S4B***), a recently described CD8 T_E_ subpopulation that constitutes an alternative MP definition with enhanced memory potential^74^.

Our basic distinction of GzmB^+^TCF1^-^ effector-type and TCF1^hi^GzmB^-^ MP P14 T_E_ populations further emphasized their substantial phenotypic heterogeneity with the latter subset displaying reduced expression of effector/activation molecules (*e.g*., CX3CR1, CXCR6, KLRD1) combined with higher levels of traditional memory markers (*e.g*., CD62L, CD127, CD27). Yet, the respective helped and helpless P14 T_E_ subsets proved strikingly similar (***Figure S4C***) with only few notable exceptions overall pointing to a subtle reduction of helpless P14 T_E_ activation (*e.g.,* universally lower expression of T-bet^56^ and of the activation-associated 130 kDa glycoform of CD43^75^, as well as a reduction of CXCR3 restricted to the TCF1^hi^GzmB^-^ compartment) (***Figures S4C-S4D***). We did not record major differences between helped and helpless P14 T_E_ regarding their LN abundance (***Figure S4E***) or functional profiles (***Figure 4K***). Interestingly, though, matching their slightly higher CD62L^hi^ MP T_E_ frequencies^74^, we detected a minor trend for higher IL-2 production *in vitro* by helpless P14 T_E_ (***Figure 4K***). Collectively, our data demonstrate a broad similarity of helped and helpless P14 T_E_ at the peak of acute LCMV infection with the exception of a subtle skewing of helpless P14 T_E_ differentiation in favor of MP rather than effector-type subsets, potentially a result of slightly delayed T_E_ trajectories. Paradoxically, helpless P14 T_E_ would therefore appear poised for better or at least commensurate P14 T_M_ formation. However, since this is apparently not their fate, these observations suggest that distinctive helpless CD8^+^ T_M_ defects are likely acquired in the post-effector phase.

### Development of helpless CD8^+^ T cell memory is contingent on the post-effector host environment

To test the above notion, we employed an *in vivo* experimental design where congenically marked helped and helpless P14 T_E_ are adoptively transferred (AT) into infection-matched helped and helpless B6 hosts on d8 after LCMV Arm infection, and P14 T_M_ formation is subsequently monitored under four distinct experimental conditions (***Figure 5A***). Supporting our previous results, absolute P14 T_M_ numbers assessed in spleens ∼7 weeks after P14 T_E_ transfers were virtually equivalent in all scenarios (***Figure 5B***). Strikingly, while both helped and helpless P14 T_E_ acquired an adequately early memory phenotype following transfer into infection-matched helped B6 recipients, the phenotypic maturation of both helped and helpless populations was significantly stunted in helpless recipients (***Figures 5C-5D***). Likewise, lower TCF1 and enhanced GzmA expression, indicating more recent cytotoxic activity/activation, strictly segregated with helpless host status but not individual – *i.e.*, helped or helpless – donor P14 T_E_ properties (***Figure 5E***). This observation was further mirrored by impaired T_M_ functionality quantified by reduced IFNγ/TNFα and IFNγ/IL-2 co-production after P14 T_E_ ATs into infection-matched helpless B6 recipients (***Figure 5F***).

**Figure 5.**
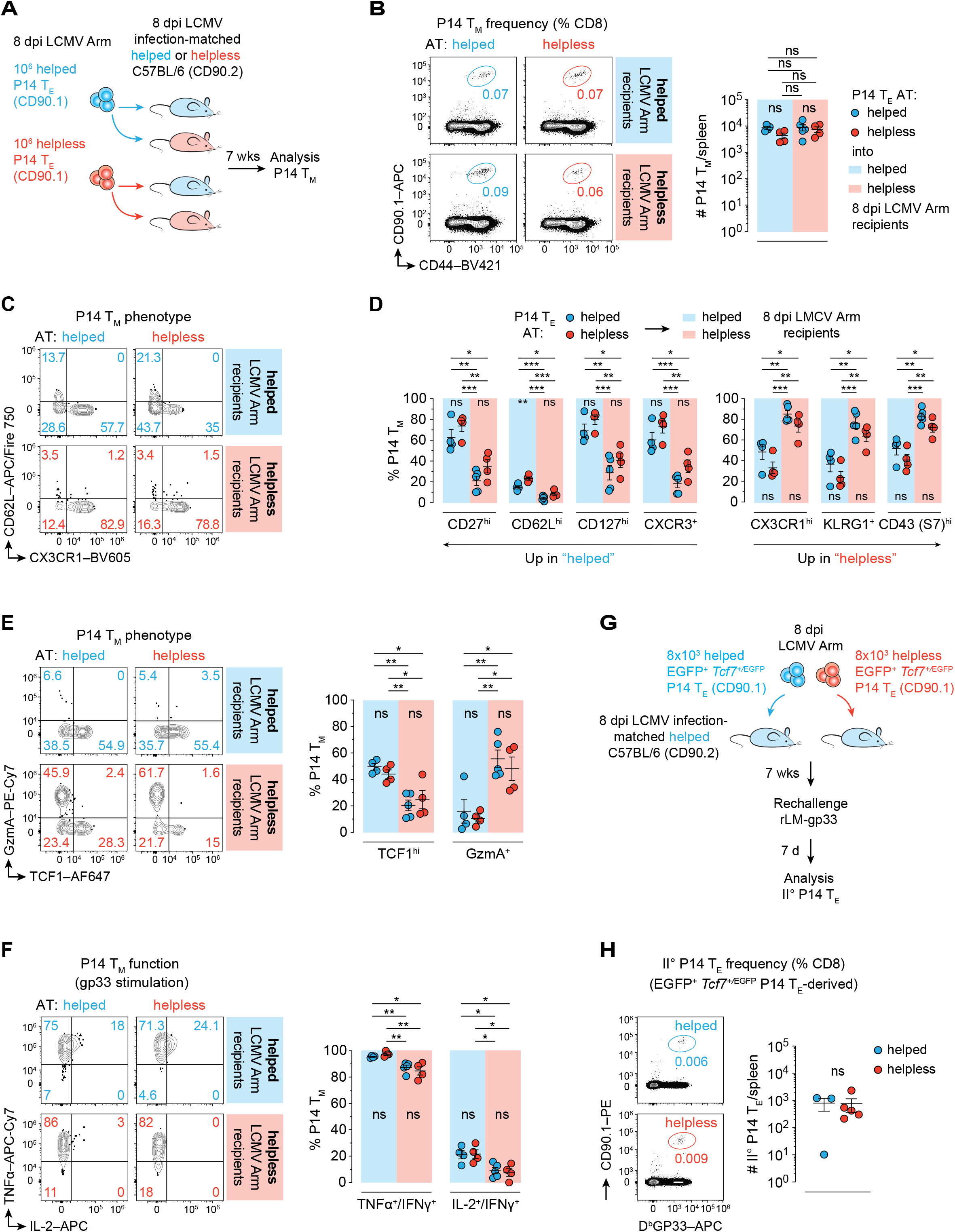
Helped *vs*. helpless post-effector environment rather than CD8^+^ T_E_ status determines CD8^+^ T_M_ fates. (**A**-**F**) Enriched helped or helpless 8 dpi P14 T_E_ were adoptively transferred into congenic infection-matched helped or helpless secondary recipients. Splenic P14 T_M_ maturation was assessed seven weeks later. (**A**) Experimental outline of P14 T_E_ “criss-cross” adoptive transfers and subsequent P14 T_M_ maturation under four distinct conditions (transient CD4^+^ T cell depletion model). (**B**) (Left) representative flow cytometry plots of frequencies and (right) enumeration of adoptively transferred P14 populations derived from helped (blue circles) or helpless (red circles) P14 T_E_ donors after maturation in respective infection-matched helped (light blue background) or helpless (light red background) secondary recipients. (**C**-**F**) Representative flow cytometry plots and summary analyses of (**C**-**D**) P14 T_M_ phenotypes, (**E**) transcription factor and effector molecule expression, and (**F**) functional properties after gp33-peptide restimulation *in vitro*. (**G**-**H**) (**G**) 8×10^3^ sort-purified 8 dpi helped or helpless EGFP^+^ *Tcf7*^+/EGFP^ P14 T_E_ were transferred into infection-matched helped secondary recipients. P14 T_M_ generation/secondary reactivity was assessed 7 weeks later after heterologous high-dose rLM-gp33 rechallenge (see ***Figure S5A***). (**H**) (Left) representative flow cytometry plots of splenic II° P14 T_E_ response to rLM-gp33 and (right) enumeration. Data representative of one out of two independent experiments with n=3-5 mice/group. Graphs plot individual helped (blue) and helpless (red) mice and mean±SEM. \**p*<0.05, \*\**p*<0.01, and \*\*\**p*<0.001 by one-way ANOVA with Tukey’s multiple comparisons test in (**A**)-(**F**) and Mann Whitney U test in (**H**). ns, non-significant.

Next, to specifically assess the memory potential of helped *vs.* helpless MP T_E_, we employed P14 chimeras generated with congenic P14 cells expressing a *Tcf7*^+/EGFP^ reporter as a proxy for intracellular TCF1 expression^76^. Notably, this reporter is different from the one used by Pais Ferreira *et al.*^72^, but reliably phenocopied our previous flow-cytometric protein analyses including higher frequencies of EGFP^+^ TCF1-expressing helpless P14 MP T_E_ in acutely-resolving LCMV Arm (***Figure S5A***). At 8 dpi, equally small numbers (8×10^3^) of sort-purified EGFP^+^ *Tcf7*^+/EGFP^ P14 MP T_E_ from either helped or helpless chimeric mice were transferred into congenic infection-matched B6 recipients to allow for subsequent P14 T_M_ formation in helped host environments (**Figure 5G**). Since direct *ex vivo* P14 T_M_ analyses in this setting are complicated by the small size of the resulting P14 T_M_ pool, B6 hosts were instead rechallenged with high-dose rLM-gp33 ∼7 weeks after the respective P14 MP T_E_ ATs, revealing commensurate II° P14 T_E_ expansions regardless of the original helped or helpless P14 T_E_ donor identities (***Figure 5H***). In parallel, we also evaluated the memory potential of EGFP^neg^ helped or helpless *Tcf7*^+/EGFP^ P14 T_E_. Here, to account for the much lower MP frequency within this population (***Figure S4A***), we opted to transfer a considerably larger number (2.5×10^5^) of helped or helpless P14 T_E_ (***Figure S5B***). Intriguingly, similar to the above observations, we also recorded an equivalent recall response by P14 T_M_ derived from EGFP^neg^ helped or helpless *Tcf7*^+/EGFP^ P14 T_E_ (***Figure S5C***). However, considering the numerical differences of adoptively transferred P14 T_E_ between these experiments together with the respective magnitude of the observed II° P14 T_E_ expansions, we estimate that both helped and helpless EGFP^+^ *Tcf7*^+/EGFP^ P14 MP T_E_ have an ∼8-fold higher memory potential than their corresponding EGFP^neg^ subsets (***Figure S5D***). Together, these observations emphasize that both helped and helpless MPs are not completely captured by a simple phenotypic signature but rather are allocated across the same “probability spectrum” of P14 T_E_ subsets with differential memory potential.

Importantly, our experiments demonstrate that the “helpless memory” fate dose not stem from diverted P14 T_E_ differentiation but rather emerges subsequently as a result of exposure to the unique helpless host environment, including potentially altered inflammatory conditions and/or prolonged viral/antigen persistence. Consistent with this contention, we observed that transfer of helped and helpless P14 T_E_ into respective naïve congenic B6 recipients, *i.e*., in the deliberate absence of host viral infection and/or inflammation, allowed for expedient and comprehensive P14 T_M_ maturation in all four experimental groups (***Figures S5E-S5J***).

### Prolonged viral antigen presentation in helpless mice

Earlier studies using transient CD4^+^ T cell depletion or constitutively CD4^+^ T cell-deficient (CD4^-/-^) mice demonstrated comparable virus control after acute LCMV in helped and helpless settings with d8 infectious titers in serum, spleen, and liver below detection threshold of conventional plaque assays^40–44^. Likewise, clearance kinetics of LCMV RNA reportedly did not differ between helped and transiently CD4^+^ T cell-depleted helpless B6 mice, generally becoming undetectable in the spleen by ∼4 weeks post infection^77^. Nevertheless, increased Ki-67 levels in early memory-phase helpless P14 T_M_ (***Figure 1F***), indicative of a recent history of activation and proliferation, suggested an antigen-driven component as a more likely explanation. We therefore sought to employ a highly sensitive bioassay that directly reports the presentation of viral antigen by proliferation of responder cells *in vivo*^78^. To this end, congenic P14 (CD45.1^+^) T_N_ were labeled with the proliferation dye CellTrace Violet (CTV), transferred into helped and helpless P14 (CD90.1^+^) chimeras at various time points post LCMV Arm, and retrieved from recipient spleens seven days later in a time course series spanning a total of eight one-week intervals from 8-383 dpi (***Figure 6A***). Despite some variability across individual helped recipients, CTV dilution by transferred responder P14 cells declined after 8 dpi and near-complete absence of measurable proliferation beyond 20 dpi indicated an effective shutdown of viral antigen presentation in these mice (***Figures 6B-6C***). In contrast, we observed substantial P14 (CD45.1^+^) responder proliferation in helpless recipients for at least two months and for up to one year after acute LCMV in individual helpless mice (***Figures 6B-6C***). Furthermore, in helpless hosts, the extent of P14 (CD45.1^+^) responder proliferation directly correlated with the phenotypic maturation status of the originally generated P14 T_M_ (CD90.1^+^) such that a particular preponderance of “immature” P14 (CD90.1^+^) T_M_ phenotypes (*i.e*., lower CD27, CD62L, CD127, CXCR3 yet increased CX3CR1, KLRG1, and CD43 expression) in some helpless mice associated with especially vigorous P14 (CD45.1^+^) responder proliferation (***Figures 6D-6E***).

**Figure 6.**
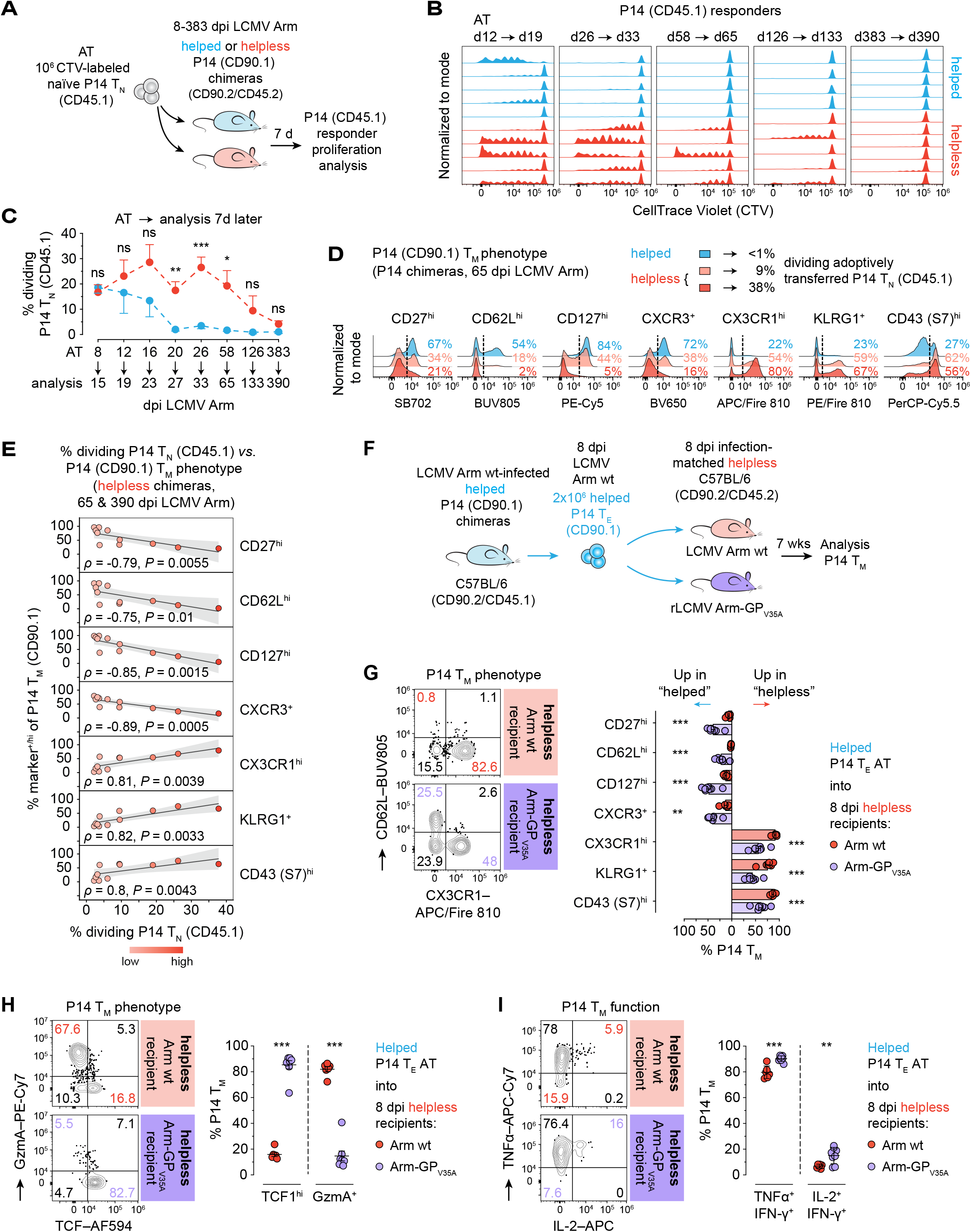
Helpless memory is sustained by prolonged cognate antigen presentation. (**A**-**E**) CellTrace Violet (CTV)-labeled P14 CD45.1^+^ T_N_ were transferred into LCMV-immune helped or helpless P14 CD90.1^+^ chimeras at various time points post LCMV Arm, recovered from spleens 7 days later, and P14 CD45.1^+^ T_N_ proliferation was quantified by CTV dilution. (**A**) Experimental setup for P14 T_N_ responder bioassay (transient CD4^+^ T cell depletion model). (**B**-**C**) (**B**) Representative histograms and (**C**) summary data of P14 CD45.1^+^ T_N_ responder proliferation over indicated weekly intervals in LCMV-immune helped and helpless P14 CD90.1^+^ chimeras. (**D**-**E**) (**D**) Representative histogram overlays of helped (blue) and helpless P14 CD90.1^+^ T_M_ featuring moderate (light red) or more pronounced immature (dark red) phenotypes and (**E**) correlation with concurrent extent of P14 CD45.1^+^ T_N_ responder proliferation. (**F**-**I**) 8 dpi helped P14 CD90.1^+^ T_E_ generated in CD45.1^+^ primary hosts were adoptively transferred into 8 dpi helpless CD90.2^+^/CD45.2^+^ secondary recipients infected with original LCMV Arm (wt; red) or a mutated gp33-determinant LCMV Arm strain (rLCMV Arm-GP_V35A_; purple) not recognized by P14 cells (see ***Figures S7A***-***S7C***). (**F**) Experimental design for P14 T_M_ maturation in helpless hosts in presence (red) or absence (purple) of cognate antigen presentation (transient CD4^+^ T cell depletion model). (**G**-**I**) Representative flow cytometry plots and summary analyses of (**G**) P14 T_M_ phenotypes, (**H**) transcription factor and effector molecule expression, and (**I**) functional properties after gp33-peptide restimulation *in vitro* as a function of helpless post-effector environments with (red) or without (purple) cognate antigen presentation. Graph in (**C**) mean±SEM of independent time course experiments with n=4-6 mice/group/time point. Spearman correlation graphs with linear regression lines in (**E**) pooled data of individual helpless P14 chimeras at 65 and 390 dpi with grey ribbons indicating 95% confidence intervals. Data in (**G**)-(**I**) representative of one out of two independent experiments with n=5-7 mice/group/experiment. \**p*<0.05, \*\**p*<0.01, and \*\*\**p*<0.001 by unpaired Student’s *t* test or Mann Whitney U test. ns, non-significant.

To formally demonstrate that viral antigen can also be recognized by CD8^+^ T_M_ under helpless conditions, we co-transferred CTV-labeled naïve P14 T_N_ (CD90.1^+^) together with P14 T_M_ (CD45.1^+^) into helped and helpless LCMV-immune congenic B6 mice at 26 dpi, followed by retrieval seven days later (***Figure S6A***). As expected, antigen-driven proliferation was almost exclusively restricted to helpless recipients for either of the transferred P14 populations (***Figure S6B***). However, in agreement with previous reports^28,79,80^, at the population level P14 T_M_ responders displayed a relatively lower sensitivity for residual LCMV antigen when compared to their naïve counterparts, and measurable proliferation was limited to a small subset that had fully diluted CTV while simultaneously upregulating Ki-67 at the end of the 7-day bioassay (***Figure S6B***). In contrast, P14 T_N_ responders proliferated more than double, yet at a comparatively slower rate than the respective P14 T_M_ subset, resulting in a ∼2-3-fold outgrowth of P14 T_M_ by P14 T_N_ in helpless recipients (***Figure S6C***). Interestingly, however, even though P14 T_N_ responders gradually acquired an antigen-experienced CD44^hi^ phenotype, they failed to downregulate CD62L suggesting a blunted/suboptimal form of CD8^+^ T cell activation reminiscent of T_N_ priming in established chronic LCMV infection^81^.

In summary, prolonged viral antigen presentation in helpless mice may permit intermittent virus-specific CD8^+^ T_M_ engagement and prompt an associated delay of timely CD8^+^ T_M_ maturation. Since the extent and frequency of such interactions likely wane with eventual antigen elimination over time, different phenotypic P14 T_M_ maturation kinetics observed in individual helpless mice (***Figure 2C***) may reflect a differential abundance of residual antigen depots capable of engaging CD8^+^ T_M_.

### CD8^+^ T_M_ maturation in the permanent absence of CD4^+^ T cell help

Our employment of the transient CD4^+^ T cell depletion model raises two important caveats: first, CD4^+^ T cells eventually repopulate the previously helpless LCMV-immune hosts (***Figure S1B***) and may even contain a virus-specific contingent primed by residual antigen^77^ that could conceivably contribute to helpless CD8^+^ T_M_ maturation; second, early studies of pathogen-specific CD8^+^ T cell memory in MHC-II-deficient (MHC-II^-/-^) mice demonstrated its progressive erosion thereby suggesting a need for CD4^+^ T cells to assure CD8^+^ T_M_ survival^12,82^. However, while a subsequent report confirmed the gradual loss of CD8^+^ T_M_ after transfer into MHC-II^-/-^ hosts, these cells were maintained in CD4^-/-^ mice suggesting that MHC-II molecules but not CD4^+^ T cells provide the relevant cues for CD8^+^ T_M_ survival^83^. We therefore sought to evaluate helpless CD8^+^ T_M_ formation, maintenance, and long-term maturation as well as the duration of cognate antigen presentation under conditions of irreversible CD4^+^ T cell deficiency (***Figure S6D***).

To this end, we compared the fate of peripheral blood LCMV-specific D^b^Gp_33_^+^ CD8^+^ T cells in CD4^-/-^ *vs*. B6 wildtype (wt) mice for ∼1.5 years after LCMV Arm infection spanning effector, early, and late memory phases (***Figure S6E***). Here, a moderate but statistically significant reduction of helpless D^b^Gp_33_^+^ CD8^+^ T_E_ (***Figure S6E***) contrasted with our previous findings in transiently CD4^+^ T cell-depleted P14 chimeras (***Figures 2B and 4B***) and this was further reflected in lower numbers of post-contraction early CD8^+^ T_M_. Moreover, in individual CD4^-/-^ mice D^b^Gp_33_^+^ CD8^+^ T_M_ continued to decline with time as indicated by widening 95% confidence intervals, which inversely correlated with subtle but progressive PD-1 re-expression suggesting potential cellular reactivation following residual antigen encounters in these mice (***Figure S6F***). Nevertheless, the evolution of helpless CD8^+^ T_M_ in CD4^-/-^ mice largely phenocopied the transient CD4^+^ T cell depletion model including delayed maturation kinetics, more pronounced phenotypic variance, and eventual acquisition of a “helped-like” mature T_M_ phenotype by ∼1.5 years after infection (***Figure S6G and cf. Figures 2C and S2H***).

Comparable timelines and trajectories were also recorded for helpless P14 T_M_ in LCMV-immune P14 chimeras generated with CD4^-/-^ recipients again highlighting a greater variability of helpless phenotypic T_M_ maturation (***Figure S6H***). Notably, however, when we compared recall responses to LCMV Arm by both “mature-like” and “immature-phenotype” helpless P14 T_M_ with their helped counterparts at ∼300 dpi, we detected near equivalent II° P14 T_E_ expansions irrespective of the original donor population suggesting that complete phenotypical T_M_ maturation also trails behind an eventual functional equilibration in CD4^-/-^ mice (***Figure S6I***) as observed after transient CD4^+^ T cell depletion (***Figures S2B and S2C***). Additionally, corresponding to the broader phenotypical variance of late-stage helpless P14 T_M_, we found considerably prolonged residual antigen presentation in individual CD4^-/-^ mice quantified by P14 T_N_ responder proliferation for >1.5 years after LCMV Arm even though the extent of proliferation in the respective mice was overall moderate and subsided by ∼2 years post infection (***Figure S6J***).

Importantly, and notwithstanding apparent differences due to the permanent lack of CD4 T cell help in acutely-resolving LCMV infection (*i.e*., moderately reduced T_E_ expansion and T_M_ survival – the latter potentially a result of enhanced cellular reactivation in response to residual antigen), these data demonstrate that helpless CD8^+^ T_M_ eventually undergo protracted but complete T_M_ maturation even under conditions of permanent CD4^+^ T cell deficiency confirming that helpless memory is in principle a reversible defect.

### Prolonged cognate antigen presentation is necessary and sufficient to impart helpless CD8^+^ T_M_ defects

The correlation of prolonged antigen presentation with the development of helpless CD8^+^ T cell memory does not exclude the possibility that altered inflammatory conditions in the helpless environment exert a supportive or even essential role during delayed CD8^+^ T_M_ maturation. To address this question, we engineered a novel recombinant LCMV Arm strain in which the valine (V) at position 35 of the LCMV glycoprotein is replaced by alanine (A) (rLCMV-GP_V35A_; ***Figure S7A***), thereby abolishing recognition by both TCR-transgenic and endogenous D^b^GP_33-41_-specific CD8^+^ T cells as previously reported for an *in vivo*-selected LCMV variant^84^. Challenge of P14 chimeric mice with rLCMV-GP_V35A_ failed to engage P14 cells or to generate an endogenous D^b^GP_33_^+^ CD8^+^ T cell response but resulted in a compensatory increase of D^b^NP_396_^+^ and D^b^GP ^+^ among CD8^+^ T_E_ populations sufficient for effective control of infectious virus (***Figures S7B-S7C***).

To specifically evaluate CD8^+^ T_M_ formation in the absence of prolonged antigen presentation but in an otherwise helpless post-LCMV inflammatory environment, we transferred helped d8 congenic P14 T_E_ (CD90.1^+^), generated in LCMV Arm wt-infected P14 chimeric mice (CD45.1^+^), into helpless B6 (CD45.2^+^) recipients that had been infected 8 days earlier with either LCMV Arm wt or rLCMV-GP_V35A_, and assessed the respective P14 T_M_ properties ∼7-9 weeks later (***Figure 6F***). In agreement with our earlier experiments, we observed that P14 T_M_ maturation was curtailed following transfer of helped P14 T_E_ into helpless LCMV Arm wt-infected secondary hosts. In contrast, the helpless milieu in rLCMV-GP_V35A_-infected recipients specifically lacking prolonged D^b^GP_33_ presentation prevented acquisition of any helpless defects and allowed for expedient phenotypical as well as functional P14 T_M_ maturation (***Figures 6G-6I***). Importantly, rLCMV-GP_V35A_-challenged hosts lacked endogenous D^b^GP_33_^+^ CD8^+^ T_M_ – ruling out a transfer of residual LCMV Arm wt virus/antigen and hence the possibility of new priming events – but continued to display higher D^b^NP_396_^+^ CD8^+^ T_M_ frequencies in an apparent echo of the I° CD8^+^ T_E_ response (***Figure S7D***). Interestingly, splenic P14 T_M_ numbers in helpless rLCMV-GP_V35A_-infected hosts were lower than those of their LCMV Arm wt-challenged recipients (***Figure S7E***), perhaps due to prolonged cognate antigen-driven proliferative expansion in the latter helpless environment (***see Figure 1F***). We also monitored the fate of helped D^b^NP_396_^+^ CD8^+^ T_E_ that were co-transferred into helpless LCMV Arm wt- or rLCMV-GP_V35A_-infected congenic recipients, *i.e.*, into a post-effector environment that permits prolonged D^b^NP_396_ presentation independent of the LCMV-GP_V35A_ mutation (**Figure S7F**). Here, we not only recovered similar numbers of D^b^NP_396_^+^ CD8^+^ T_M_ from spleens of both recipient groups (***Figure S7E***) but observed the very same extent of helpless CD8^+^ T_M_ features including low TCF1 and high GzmA expression (***Figure S7F***).

Collectively, our data illustrate that prolonged exposure of post-effector CD8^+^ T cells to their cognate antigen is a key “driver” for the development of helpless memory. Although we did not directly quantify inflammatory parameters in the above experiments, the particular constellation of shared and diverging fates of P14 T_M_ and D^b^NP_396_^+^ CD8^+^ T_M_ in LCMV Arm wt-*vs.* rLCMV-GP_V35A_-infected hosts permits the conclusion that prolonged cognate antigen presentation under helpless conditions is both a necessary and sufficient condition to promote the development of helpless CD8^+^ T cell memory.

## DISCUSSION

Here, we longitudinally assessed the fate of specific CD8^+^ T cells after acutely-resolving LCMV infection in the absence of CD4^+^ T cell help and identify sustained cognate antigen presentation as the underlying cause for their signature memory defect, *i.e.*, poor recall expansion^10–12^. Mechanistically, our data demonstrate that helpless CD8^+^ T cell memory is not “imprinted” during CD8^+^ T_E_ differentiation but emerges as a consequence of developing and established CD8^+^ T_M_ exposure to the post-effector helpless environment resulting in delayed but not arrested T_M_ maturation. Importantly, as cognate antigen presentation is effectively eliminated over time, helpless CD8^+^ T_M_ continue to evolve and eventually become phenotypically and functionally indistinguishable from their helped counterparts. Notwithstanding kinetic differences, these observations applied to both TCR-transgenic and endogenously generated LCMV-specific CD8^+^ T_M_ in transient as well as permanently CD4-deficient settings. Redefining helpless CD8^+^ T cell memory as a temporary condition, our study proposes a unifying explanation for this phenomenon with general implications for our understanding of CD8^+^ T_M_ development.

Failure to timely eliminate antigen after infection may result in context-specific adaptations to CD8^+^ T_M_ formation, including clonal deletion and cellular exhaustion in chronic viral infections (*e.g.*, LCMV cl13)^59,85^ or “memory inflation”, a result of periodic restimulation, in non-productive viral latency (*e.g*., cytomegalovirus)^86^. However, local residual antigen reservoirs in acutely-resolving viral infections (*e.g.*, influenza) have also been shown to sustain or even improve cellular immunity, presumably by promoting a more effector-like CD8^+^ T_M_ state^87^, while lingering antigen after SARS-CoV-2 infection, *e.g.* in long COVID, may exacerbate pathology in part by precluding acquisition of protective CD8^+^ T_M_ potential^88,89^. In the helpless settings interrogated here, we demonstrate a dynamic immune response equilibrium that emphasizes the sensitivity of developing CD8^+^ T_M_ to prolonged antigen presentation, a likely role for stochastic antigen encounters and their impact on T_M_ evolution across individual mice, as well as a notably extended time course until full systemic helpless T_M_ maturation is achieved. However, unlike the permanent scarring of “recovering” exhausted T_EX_ after chronic viral infection^90^, our model rather constitutes a threshold scenario in which transient antigen persistence leaves a defined but temporary defect in the evolving CD8^+^ T_M_ pool. Importantly, system-specific variables (*e.g.,* nature, route, and dosage of antigen challenge; mode and duration of CD4^+^ T cell exclusion from the immune response; scope, detail, and timing of analytical readouts) may thus differentially shape the fate of long-term helpless CD8^+^ T cell memory and the concept of prolonged antigen presentation in specific models will need to be experimentally validated.

Our conclusions seemingly differ from the invocation of CD8^+^ T cell “exhaustion” in an experimental system of helpless CD8^+^ T_M_ generation by DNA vaccination^22,65,91^. In this setup, helpless vaccine-specific CD8^+^ T_E_ were less frequent and characterized by numerous molecular, phenotypic, and functional alterations that appear to carry forward into a smaller helpless CD8^+^ T_M_ pool. Given the complexities of this essentially CD4^+^ T cell help-dependent model, it is difficult to predict how prolonged antigen presentation, not yet experimentally interrogated, may compound CD8^+^ T_M_ maturation. However, the extent to which this or other helpless models may promote genuine CD8^+^ T cell exhaustion is contingent on particular experimental constraints, a clarification of the role of system-specific antigen persistence, and ultimately the chosen definition for CD8^+^ T cell exhaustion^85,92^. In our models of acutely-resolving helpless LCMV infection, we have not obtained any evidence for robust exhaustion signatures. Nevertheless, one of the key conclusions drawn from the DNA vaccination model, namely that lack of CD4^+^ T cell help may impair CD8^+^ cytotoxic differentiation^65^ is in fact echoed by our observation of a minor numerical reduction of the activated helpless P14 T_E_ compartment.

Our studies with a mutant LCMV strain not recognized by P14 cells imply a likely negligible role for altered inflammatory conditions in the pathogenesis of helpless CD8^+^ T cell memory. Similarly, notwithstanding subtle differences, robust transcriptional, phenotypic, and functional effector profiles as well as commensurate memory potential by helpless CD8^+^ T_E_ make altered CD8^+^ T cell immunity an unlikely candidate for the failure of timely antigen removal in this experimental setup. In contrast, minor alterations and a somewhat protracted course, *e.g.*, progressive PD-1 re-expression as well as an associated gradual loss of specific CD8^+^ T_M_, in individual CD4^-/-^ mice precluding virus-specific “late-comers” from a resurging CD4^+^ T cell compartment^77^ point to direct CD4^+^ T cell-mediated activities that provide the necessary contributions for efficient residual antigen elimination. Future work will thus need to identify the cellular source(s) of prolonged antigen presentation, and to detail how different CD4^+^ T cell subsets (including CD4^+^ regulatory T cells operating the immediate post-effector phase vial IL-10 and cytotoxic T-lymphocyte associated protein 4/CTLA-4^93,94^), relevant effector mechanisms and cellular interaction partners (most prominently, arguably, B cells^95^) are engaged across different anatomic niches and time (*i.e*., priming, resolution, maintenance) to promote an expedient suppression of antigen presentation.

In the meantime, our observations may serve to situate different shades of “helplessness” across distinct experimental or naturally occurring scenarios, and longitudinal assessment of circulating CD8^+^ T_M_ may present a valuable tool both reporting past antigen exposure and providing an outlook of future maturation potential. Furthermore, these data suggest that optimized immunization strategies seeking to boost adaptive immune responses will need to consider prolonged antigen presentation as a potential caveat limiting acquisition of protective CD8^+^ T_M_ properties that may be mitigated by engagement of adaptive immune response across both the CD8^+^ and CD4^+^ T cell compartments in addition to specific B cell immunity. Finally, identifying potential pathways that may compensate for a lack of CD4^+^ T cell help during priming may benefit the development of improved therapeutic avenues, especially in immunocompromised individuals or settings of cancer.

### Limitations of the study

Although readily integrated into a longitudinal model of dynamic CD8^+^ T_M_ maturation, our analyses are largely limited to circulating CD8^+^ T_M_ in a single host-pathogen system. Further studies are required to assess the scope of our conclusions (*e.g.*, for resident memory CD8^+^ T cell populations) in additional model systems for acute pathogen infection as well as vaccination, autoimmune and/or cancer models, and to assess their general applicability to human CD8^+^ T cell responses. Moreover, how specifically, *i.e.*, by which molecular and cellular interactions, CD4^+^ T cell help shapes the outcome of CD8^+^ T cell responses to prevent helpless memory defects in these settings will need to be addressed in future work. Similarly, identifying the cellular source(s) and mechanisms of cognate antigen presentation to helpless CD8^+^ T_M_ *in vivo* will provide insight for preventive and therapeutic strategies.

## ACKNOWLDEGMENTS

We thank Pei-Yu Kuo (Cytek Biosciences, USA) for expert technical assistance with cell sorting, Tom Nguyen, Kelsey Haist, and Haedar Abuirqeba for assistance with experiments, Ross Kedl (University of Colorado – Anschutz Medical Campus, Aurora, CO, USA) for critical reading of the manuscript, as well as all past and current members of the laboratories of D.H. and A.O.K. for scientific discussions. We thank Gregoire Lauvau (Albert Einstein College of Medicine, New York, NY, USA) for wildtype *Listeria monocytogenes* as well as for advice and protocols regarding bacterial propagation and murine infections. We thank the NIH Tetramer Core Facility, the ISMMS Dean’s Flow Cytometry Core, the ISMMS Human Immune Monitoring Center, and the ISMMS Center for Comparative Medicine and Surgery. This research was supported by German Research Foundation (DFG) research fellowship HE 7559/1-1 (V.v.d.H.); NIH T32 training grants AI052066 and DK007792 (B.D.); NIH grants R21AI121840, R21AI169789 and RO1AI142985 (J.C.-d.l.T.); and NIH grants R01AI093637 and R01DK130425 (D.H.).

## AUTHOR CONTRIBUTIONS

Conceptualization: V.v.d.H., D.C., R.A, and D.H.; investigation, V.v.d.H., B.D., B.C, K.J., E.H., K.A., T.D., and G.F.; formal analysis, V.v.d.H., V.R., and D.H.; software, V.R.; writing – original draft, V.v.d.H. and D.H.; writing – review & editing, all authors; visualization, V.v.d.H., V.R., and D.H.; resources, A.K., J.d.l.T., and D.H.; funding acquisition: V.v.d.H., B.D., and D.H.; supervision, D.H.

## METHODS

### RESOURCE AVAILABILITY

#### Lead Contact

Further information and requests for resources and reagents should be directed and will be fulfilled by the lead contact, Dirk Homann.

#### Materials availability

Engineered rLCMV Armstrong-GP_V35A_ used in this study is available from the lead contact upon request. No other new unique reagents were generated.

#### Data and code availability

scRNAseq data are available from the National Center for Biotechnology Information Gene Expression Omnibus under accession GSE251744. Analyses were done with custom python or R scripts, respectively, using standard python/R packages as referenced in key resources table and source code is provided upon request. Any additional information required to reanalyze the data reported in this paper is available from the lead contact upon request.

### EXPERIMENTAL MODEL AND SUBJECT DETAILS

#### Mice

C57BL/6J (B6; #000664), B6.PL-*Thy1^a^*/CyJ (B6 CD90.1^+^; #000406), B6.SJL-*Ptprc^a^ Pepc^b^*/BoyJ (B6 CD45.1^+^; #002014), B6.129S2-*Cd4^tm1Mak^*/J (CD4^-/-^; #002663), and B6(Cg)-*Tcf7^tm1Hhx^*/J mice (*Tcf7*^+/EGFP^; #030909) were purchased from The Jackson Laboratory. TCR-transgenic P14 mice specific for the dominant D^b^-restricted LCMV glycoprotein GP_33-41_-epitope were originally obtained from Michael Oldstone (The Scripps Research Institute, La Jolla, USA) and maintained on a B6 CD90.1^+^ or B6 CD45.1^+^ background. B6(Cg)-*Tcf7^tm1Hhx^*/J mice were backcrossed with P14 CD90.1^+^ mice onto the B6 background to obtain *Tcf7*^+/EGFP^ P14 CD90.1^+^ TCF1-EGFP heterozygous reporter mice preserving the T cell memory-associated p45 TCF1 isoform^96^. Both male and female mice between 6-10 weeks of age were used in age- and sex-matched experimental cohorts. Only sex-matched donor-recipient adoptive transfers were performed. All mice were housed in ventilated cages at ∼20 °C and ∼55% humidity with 12h/12h light-dark cycles under specific-pathogen-free conditions at the Icahn School of Medicine at Mount Sinai (ISMMS) vivarium. All procedures were approved by the ISMMS Institutional Animal Care and Use Committee, and all efforts were made to minimize animal suffering.

#### Cell lines

Vero E6 cells (Vero C1008, ATCC CRL-1586) were grown in Eagle’s Minimum Essential Medium (EMEM, ATCC) supplemented with 10% FBS (Gibco) and 1% penicillin/streptomycin (Gibco). BHK-21 cells (BHK-21 C-13, ATCC CCL7) were cultured in Dulbecco’s Modified Eagle Medium (DMEM, ATCC) with 10% FBS, 1% penicillin/streptomycin, and 5% tryptose phosphate broth (Sigma). Cell lines were maintained at 37°C, 5% CO_2_, and >95% humidity.

### METHOD DETAILS

#### Adoptive T cell transfers

To generate P14 chimeras, naïve helped/helpless B6 or CD4^-/-^ recipients were adoptively transferred intravenously (i.v.) with 5×10^3^ splenic P14 T_N_ cells congenic for CD90.1^+^ or CD45.1^+^ isolated by negative selection using the EasySep Mouse CD8+ T cell isolation kit (STEMCELL Technologies) before LCMV Arm infection eight hours later. For rechallenge experiments, splenic P14 T_M_ were enriched at indicated time points after LCMV Arm by removal of APC- or PE-labeled CD4^+^ T and CD19^+^ B cells with anti-APC/PE magnetic microbeads (Miltenyi Biotec & STEMCELL Technologies) and 5×10^3^ P14 T_M_ were transferred i.v. into naïve congenic hosts prior to LCMV Arm or rLM-gp33 challenge. For “criss-cross” adoptive transfer experiments, helped or helpless splenic P14 T_E_ CD90.1^+^ were enriched at 8 dpi after removal of CD4^+^ T, CD19^+^ B, and CD90.2^+^ non-P14 T cells as described above, and 1-2×10^6^ cells were transferred i.v. into congenic 8 dpi LCMV Arm or rLCMV Arm-GP_V35A_ infection-matched helped or helpless recipients as specified. In some experiments, 7.5×10^5^ D^b^NP_396_^+^ CD45.1^+^ CD8^+^ T_E_ were co-transferred along with P14 T_E_ into congenic 8 dpi LCMV Arm or rLCMV Arm-GP_V35A_ infection-matched helpless hosts. For MP T_E_ transfer experiments, EGFP^+^ and EGFP^neg^ *Tcf7*^+/EGFP^ P14 T_E_ CD90.1^+^ were sort-purified from spleens of helped or helpless *Tcf7*^+/EGFP^ P14 CD90.1^+^ chimeras generated as detailed above and 8×10^3^ EGFP^+^ or 2.5×10^5^ EGFP^neg^ P14 T_E_, respectively, were transferred i.v. into congenic 8 dpi LCMV Arm infection-matched helped recipients. To induce larger numbers of P14 T_M_ with enhanced memory maturation kinetics, 5×10^4^ P14 CD45.1^+^ T_N_ were transferred i.v. into naïve congenic B6 hosts followed by LCMV Arm and P14 T_M_ were recovered from spleens ∼30 dpi for secondary transfers into helped and helpless B6 recipients for detection of residual cognate antigen as outlined below.

#### Transient CD4^+^ T cell depletion

For helpless LCMV priming, P14 chimeras or B6 mice were transiently depleted of CD4^+^ T cells by intraperitoneal (i.p.) injection of two consecutive doses of 200 µg of anti-CD4 antibody (clone GK1.5, Biolegend) on d-1 and d+1 in relation to LCMV infection. Rat IgG2b isotype antibody (clone RTK4530, Biolegend) was injected similarly to generate helped control mice. Efficiency of CD4^+^ T cell depletion was routinely assessed in small amounts of peripheral blood on d0 prior to LCMV challenge by flow cytometry staining with a non-competitively binding anti-CD4 antibody (clone RM4-4, Biolegend).

#### Viral and bacterial Infections

LCMV strains Armstrong 53b and cl13 were originally obtained from Michael Oldstone (The Scripps Research Institute, La Jolla, USA). Recombinant rLCMV Armstrong-GP_V35A_ was engineered as detailed below. LCMV strains were grown in BHK-21 cells in DMEM with 2% FBS, and infectious viral titers were quantified by plaque assay on Vero E6 monolayers as described^97,98^. For acute infections, mice were injected i.p. with 2×10^5^ plaque forming units (PFU) LCMV Arm or rLCMV Arm-GP_V35A_. LCMV-immune mice were monitored for up to ∼2.5 years and aging mice with gross physical abnormalities (*e.g.*, severe skin lesions, emaciation), tumors, or T cell clonal expansion among virus-specific CD8^+^ T_M_ populations were excluded from the study. Lifelong chronic infection with 2×10^6^ PFU LCMV cl13 was induced by i.v. route in CD4^+^ T cell depleted mice to generate exhausted virus-specific CD8^+^ T cells. Viral load was quantified by plaque assay as previously described^97,98^. Recombinant *Listeria monocytogenes* expressing the LCMV-gp_33-41_ (rLM-gp33) determinant was originally provided by Hao Shen^11^ and grown in Brain Heart Infusion broth (BD Biosciences) to an OD_600_ ∼0.1. For rLM-gp33 challenges after adoptive P14 T_M_ transfers into naïve recipients or heterologous infections of LCMV-immune mice, 5×10^4^ colony forming units (CFU) were injected i.v.. Bacterial inoculum titers were assessed by incubating bacterial dilutions on Brain Heart Infusion agar (BD Biosciences) plates overnight.

#### Generation of recombinant rLCMV-GP_V35A_

Recombinant rLCMV-GP_V35A_ precluding recognition by TCR-transgenic and endogenous D^b^GP_33-41_-specific CD8^+^ T cells was engineered using established plasmid-based LCMV reverse genetics^99–101^. Briefly, two overlapping PCR products were amplified from pCAGGS-ARM/GPC (= glycoprotein precursor) plasmid template with the following primer pairs incorporating the mutant amino acid change V35A (Valine to Alanine at position 35; mutation highlighted in bold lower case): PCR fragment 1 (230 bp) primers 5’-GACCGGCGGCTCTAGAGCCTC-3’ and 5’-**cg**CAGCCTTGATACCCGTGATCACG-3’; PCR fragment 2 (1054 bp) primers 5’-GGTATCAAGGCTG**cg**TACAATTTTGCCACCTGTGGGATATTC-3’ and 5’-TTGCATGTTCTAGGTACCAAAACTTTGAGTAATTG-3’. The two PCR products were cloned via In-Fusion HD Cloning (Takara) into the pCAGGS-ARM/GPC plasmid digested with XbaI and KpnI replacing a segment of the wild-type GPC sequence. The resulting pCAGGS-ARM/GPC (V35A) plasmid containing the V35A mutation was confirmed by sequencing and expression in transfected cells. A 1528 bp fragment corresponding to the full-length GPC open reading frame (ORF) containing mutation V35A and 15 bp nucleotides complementary to the cloning S segment cassette was PCR-amplified from pCAGGS-ARM/GPC (V35A) using primers 5’-CCTATCCTACAGAAGGATGGGTCAGATTGTGACAATGTTTGAG-3’ and 5’ GGGAGGCGCTGTTCTTCAGCGTCTTTTCCAGACGGTTTT-3’, and in-fusion cloned (Takara) into the GPC locus of the mPol1 LCMV S NP/BsmBI cassette digested with BsmBI. The resulting plasmid mPol1/LCMV Sag (GPC V35A) was confirmed by sequencing. To rescue rLCMV-GP_V35A_, BHK-21 cells (7×10^5^/well) were transfected in tissue culture-treated 6-well plates using Lipofectamine 2000 (ThermoFisher Scientific) with the LCMV Arm plasmids pCAGGS-NP (0.8 µg), pCAGGS-L (1 µg), mPol1/LCMV Sag (GPC V35A) (0.8 µg), and mPol1/LCMV Lag (1.4 µg). Transfection medium was replaced 5h later with 3 mL of DMEM containing 2% FBS and tissue culture supernatants (TCS) collected at days 3 and 6 post transfection were titered by indirect fluorescence focus assay as described^101^. Day 6 TCS was used to infect BHK-21 monolayers (multiplicity of infection/MOI = 0.01), viral TCS was collected 72h later, titered, and fresh BHK-21 cells were again infected at a MOI = 0.01 to generate a working stock of rLCMV-GP_V35A_. RNA was isolated from virus-infected cells 48h later, the viral glycoprotein ORF was amplified by RT-PCR, and the V35A mutation was confirmed by sequencing.

#### *In vivo* P14 responder bioassay

For P14 responder assays, magnetically isolated splenic P14 CD45.1^+^ T_N_ (EasySep Mouse CD8+ T cell isolation kit, STEMCELL Technologies) were labeled with CellTrace Violet (CTV, Invitrogen) according to manufacturer’s instructions, and ∼10^6^ P14 T_N_ were injected i.v. into helped or helpless LCMV-immune congenic P14 chimeras at indicated time points. P14 cells were retrieved from recipient spleens 7d later to assess their proliferation by CTV dilution using flow cytometry. In some cases, 7.5×10^5^ CTV-labeled splenic P14 CD45.1^+^ T_M_, enriched by removal of PE-labeled CD4^+^ T and CD19^+^ B cells as detailed above, were co-transferred together with 10^6^ P14 CD90.1^+^ T_N_.

#### Mouse tissue processing

Peripheral blood was obtained via cheek bleeds, subjected to red blood cell (RBC) lysis by ACK lysis buffer (Gibco), and PBMCs were stained with antibodies for flow cytometry as described below. Naïve, effector, or memory P14 cells for functional assays or flow cytometric analyses were retrieved from spleen or lymph node single cell suspensions as specified after disrupting spleens through 70 µm filters using 3 mL-syringe plungers or meshing lymph nodes between sand-blasted ends of frosted microscopy slides. Splenic RBCs were subsequently lysed using ACK buffer. In some cases, mice were injected i.v. with 3 µg of APC-labeled anti-CD8b.2 antibody (clone 53-5.8, Biolegend) 5 min prior to euthanasia to mark vasculature-associated cells *in vivo*.

#### Flow cytometry and cell sorting

For surface staining, PBMCs, spleen, or LN single cell suspensions were incubated with fluorescently labeled antibodies or anti-LCMV D^b^-restricted tetramers (NIH Tetramer Core) as well as anti-CD16/32 (clone S17011E, Biolegend) in FACS buffer (phosphate buffered saline/PBS with 0.5% fetal bovine serum/FBS, 0.5% 3 mM sodium azide, and 0.5 mM EDTA) for 45 min at 4°C in the dark. To exclude dead cells from downstream analysis, spleen and LN samples were stained with LIVE/DEAD Fixable Blue (ThermoFisher Scientific), Zombie UV, Violet, Aqua, Green, or NIR Fixable viability dye (all Biolegend) in PBS for 10 min at RT immediately prior to fixation. PBMCs were fixed at room temperature (RT) with Cytofix reagent (BD Biosciences) and samples were subsequently washed with BD Cytoperm solution (BD Biosciences) to clear residual RBCs. Stained spleen or LN samples were fixed with 1-4% phosphate-buffered paraformaldehyde solution (FluoroFix or Fixation Buffer, Biolegend). To detect transcription factors and certain intracellular proteins, samples were fixed/permeabilized over-night at 4°C using the FoxP3 transcription buffer set (ThermoFisher Scientific). For intracellular cytokine staining after gp33 peptide stimulation, samples were fixed/permeabilized for 20 min at RT with the BD Cytofix/Cytoperm kit (BD Biosciences) as per manufacturer’s instructions. To preserve intranuclear EGFP signal before intracellular staining, samples were either fixed/permeabilized at RT for 15 min with BD Cytofix reagent (BD Biosciences) or alternatively subjected to 15 min of 2% phosphate-buffered PFA prefix^102^ followed by subsequent overnight fixation/permeabilization at 4°C with the FoxP3 transcription buffer set. Intracellular/intranuclear staining was conducted by incubation of fixed/permeabilized cells with unconjugated, biotin-conjugated, or fluorescently labeled primary antibodies in Cytoperm (BD Biosciences) or FoxP3 Transcription buffer set permeabilizing reagent for 60 min at RT. In some cases, secondary intracellular staining was performed for 25 min at RT using cross-adsorbed fluorescently labeled anti-rabbit secondary antibodies or fluorescently-conjugated streptavidin. BD Horizon Brilliant Stain Buffer Plus (BD Biosciences) was added to antibody master mixes where more than two Brilliant Ultraviolet- or Brilliant Violet-labeled antibodies were used. Samples were resuspended in FACS buffer prior to acquisition and 1X Tandem Stabilizer reagent (Biolegend) was added if samples were not immediately analyzed. All samples were acquired on a 2-laser FACSCalibur, 4-laser LSRII, 3-laser Canto II, or 5-laser LSRFortessa X-20 (all BD Biosciences), a 3-laser Attune NxT (ThermoFisher Scientific), or a 5-laser Aurora (Cytek Biosciences) flow cytometer. Data were analyzed with FlowJo version 10 (BD Biosciences). To assess P14 T_N_ responder proliferation, the FlowJo proliferation platform was used to calculate “% dividing cells”, *i.e.*, the cellular precursor frequency ranging from 0 to 1 indicating the probability that a cell will divide at least once^103^. For adoptive co-transfer experiments with limited proliferation or low cell numbers, the fraction of CTV diluted was directly quantified instead by gating on all P14 T_N_ or T_M_ that divided at least once. Heatmap visualization of row-normalized gMFI values was generated using the ComplexHeatmap package (v 2.14.0) in R. For scRNAseq experiments, live P14 CD90.1^+^ T_E_ (LIVE/DEAD Blue^neg^ CD8a^+^ CD90.1^+^ CD4^neg^ CD19^neg^) from spleens of helped and helpless P14 chimeras at 7 dpi were sorted on a 5-laser Aurora CS sorter (Cytek Biosciences). For P14 T_E_ subset sorts on d8, live EGFP^+^ or EGFP^neg^ *Tcf7*^+/EGFP^ P14 CD90.1^+^ T_E_ (Zombie Aqua^neg^ EGFP^+^ or EGFP^neg^ CD8a^+^ CD90.1^+^) from helped or helpless spleens were sorted on a 5-laser FACS Aria II sorter (BD Biosciences). Small aliquots of sorted samples were checked for purity (on average >98%). Antibodies and reagents were purchased from Biolegend, BD Biosciences, ThermoFisher Scientific, Miltenyi Biotec, Cell Signaling, R&D Systems, Jackson ImmunoResearch, and further details are found in key resources table.

#### T cell stimulation and cytokine staining

To assess cytokine production, spleen single cell suspensions were incubated for 4h at 37°C in the presence of gp33 peptide (1 µg/mL KAVYNFATC; Genscript), brefeldin A (5 µg/mL; Biolegend), and monensin (2 µM; Biolegend). Phorbol 12-myristate13-acetate (PMA)/ionomycin-stimulated (Cell Activation cocktail, Biolegend) and unstimulated samples in the absence of peptide were included as positive and negative controls, respectively. In some cases, samples were pre-treated for 1h at 37°C with 0.1 mM TAPI-2^104^ (R&D Systems) prior to subsequent incubation with gp33 peptide as detailed above in the continuous presence of TAPI-2. At the end of the stimulation period, cultures were washed in FACS buffer and kept at 4°C until flow cytometry staining.

#### Single-cell RNA sequencing (scRNAseq)

##### Cell processing, library preparation, and sequencing

Splenic P14 T_E_ were recovered from helped or helpless P14 CD90.1^+^ chimeras 7 dpi with LCMV Arm, processed and sort-purified as detailed above. Viability and cell numbers were determined by ViaStain acridine orange/propidium iodide staining (Nexelcom) and average viability across all samples (n=4 replicates/condition) was >94% (range: 87.4-97.4%). scRNAseq was performed using the Chromium platform (10x Genomics) with the 5’ gene expression (5’ GEX) v2 kit according to manufacturer’s instructions and a targeted recovery of 8×10^3^ cells/sample. Briefly,

Gel-Bead in Emulsions (GEMs) were generated on the sample chip in a Chromium X/iX controller. Barcoded cDNA was extracted from the GEMs by Post-GEM RT-cleanup and amplified for 13 cycles. Amplified cDNA was fragmented and subjected to end-repair, poly A-tailing, adapter ligation, and 10x-specific sample indexing following the manufacturer’s protocols. Libraries were quantified using QuBit (ThermoFisher Scientific) and TapeStation (Agilent Technologies) analyses and were sequenced in pair-end mode on a NovaSeq instrument (Illumina) targeting a depth of 25,000 reads per cell.

#### scRNAseq preprocessing and analysis

Fastq files were generated and aligned to reference transcriptome mm10-2020-A using the Cell Ranger pipeline version 5.0.0 (10x Genomics)^105^. Downstream analysis was performed in Python (version 3.10.8) using the Scanpy toolkit (version 1.9.3)^106^. Briefly, obtained gene-level gene expression data were filtered (only cells with: >200 but <2,500 genes/cell, <25% mitochondrial genes/cell, only genes found in >3 cells), total count-normalized, log1p transformed and highly variable genes were extracted. Filtered data were subjected to PCA dimensionality reduction and projected into 2D UMAP embedding. Unsupervised Leiden-graph clustering^67^ was performed with resolution parameter set at 0.4. Marker genes were identified by *t*-test with overestimated variance “one-versus-everyone” and ranked according to fold change. Sample-biased cluster “C5” (see ***Figure S3B***, driven by sample “H3”) was excluded from downstream analyses. Clusters were annotated based on marker gene expression profiles, gene set enrichment analyses using ClusterProfiler (version 3.8, R v 4.2.2), and after projection of a reference dataset^68^ using the SingleR (version 3.18) method^107^. Cell type quantifications and statistics as well as graphical plotting were performed in Python/Scanpy using Matplotlib (version 3.7.2) and pandas package (version 1.5.2) functionalities. *Z* score data of log2-normalized gene expression profiles across clusters were additionally visualized using the ComplexHeatmap package (version 2.14.0) in R. Differential gene expression within each cluster between conditions (“helped” *vs.* “helpless”) was conducted by pseudo-bulk analysis using the Python decoupleR package (version 1.5.0)^108^. Differential expression analyses were visualized using the EnhancedVolcano package (version 1.13.2) in R.

## QUANTIFICATION AND STATISTICAL ANALYSIS

Data analysis and graphical representation were performed in GraphPad Prism 9 and 10 (GraphPad Software) or R (version 4.2.3) and edited for appearance using Adobe Illustrator (Adobe). Normal distribution was determined by Shapiro Wilk test and visualization by quantile-quantile plots. Statistical significance was assessed by unpaired Student’s *t* tests or non-parametric Mann-Whitney U test for comparison between two groups, as well as by one-way ANOVA with Tukey’s (comparing every mean with every other mean) or Šídák’s (comparing selected pairs of means) multiple comparisons post hoc testing or repeated-measures one way ANOVA with Šídák’s multiple comparisons post hoc testing for analysis of more than two groups with normally distributed values as specified in ***Figure Legends***. Spearman’s rank correlation was computed for non-parametric associations between two variables and visualized in GraphPad Prism 10 or using the ggpubr R package (version 0.6.0). Principal component analyses and biplots were computed and visualized in R using the FactoMineR (version 2.8) and factoextra (version 1.0.7) packages. Any additional R packages employed for data handling and graphical display are listed in key resources table. Data are displayed as mean±SEM or 95% confidence intervals unless indicated otherwise adopting the following convention: \**p* < 0.05, \*\**p* < 0.01, \*\*\**p* < 0.001, ns, non-significant.

## REFERENCES

1. Williams, M.A., and Bevan, M.J. (2007). Effector and memory CTL differentiation. Annu Rev Immunol 25, 171–192. 10.1146/annurev.immunol.25.022106.141548.

2. Khanolkar, A., Badovinac, V.P., and Harty, J.T. (2007). CD8 T cell memory development: CD4 T cell help is appreciated. Immunol Res 39, 94–104. 10.1007/s12026-007-0081-4.

3. Wiesel, M., and Oxenius, A. (2012). From crucial to negligible: functional CD8(+) T-cell responses and their dependence on CD4(+) T-cell help. Eur J Immunol 42, 1080–1088. 10.1002/eji.201142205.

4. Laidlaw, B.J., Craft, J.E., and Kaech, S.M. (2016). The multifaceted role of CD4(+) T cells in CD8(+) T cell memory. Nat Rev Immunol 16, 102–111. 10.1038/nri.2015.10.

5. Borst, J., Ahrends, T., Babala, N., Melief, C.J.M., and Kastenmuller, W. (2018). CD4(+) T cell help in cancer immunology and immunotherapy. Nat Rev Immunol 18, 635–647. 10.1038/s41577-018-0044-0.

6. Wu, R., and Murphy, K.M. (2022). DCs at the center of help: Origins and evolution of the three-cell-type hypothesis. J Exp Med 219. 10.1084/jem.20211519.

7. Bachmann, M.F., Zinkernagel, R.M., and Oxenius, A. (1998). Immune responses in the absence of costimulation: viruses know the trick. J Immunol 161, 5791–5794.

8. Le Bon, A., Etchart, N., Rossmann, C., Ashton, M., Hou, S., Gewert, D., Borrow, P., and Tough, D.F. (2003). Cross-priming of CD8+ T cells stimulated by virus-induced type I interferon. Nat Immunol 4, 1009–1015. 10.1038/ni978.

9. Bevan, M.J. (2004). Helping the CD8(+) T-cell response. Nat Rev Immunol 4, 595–602.

10. Janssen, E.M., Lemmens, E.E., Wolfe, T., Christen, U., von Herrath, M.G., and Schoenberger, S.P. (2003). CD4+ T cells are required for secondary expansion and memory in CD8+ T lymphocytes. Nature 421, 852–856.

11. Shedlock, D.J., and Shen, H. (2003). Requirement for CD4 T cell help in generating functional CD8 T cell memory. Science 300, 337–339.

12. Sun, J.C., and Bevan, M.J. (2003). Defective CD8 T cell memory following acute infection without CD4 T cell help. Science 300, 339–342.

13. Harty, J.T., and Badovinac, V.P. (2008). Shaping and reshaping CD8+ T-cell memory. Nat Rev Immunol 8, 107–119. nri2251 [pii] 10.1038/nri2251.

14. Gressier, E., Schulte-Schrepping, J., Petrov, L., Brumhard, S., Stubbemann, P., Hiller, A., Obermayer, B., Spitzer, J., Kostevc, T., Whitney, P.G., et al. (2023). CD4(+) T cell calibration of antigen-presenting cells optimizes antiviral CD8(+) T cell immunity. Nat Immunol 24, 979–990. 10.1038/s41590-023-01517-x.

15. Intlekofer, A.M., Takemoto, N., Kao, C., Banerjee, A., Schambach, F., Northrop, J.K., Shen, H., Wherry, E.J., and Reiner, S.L. (2007). Requirement for T-bet in the aberrant differentiation of unhelped memory CD8+ T cells. J Exp Med 204, 2015–2021. jem.20070841 [pii] 10.1084/jem.20070841.

16. Cox, M.A., and Zajac, A.J. (2010). Shaping successful and unsuccessful CD8 T cell responses following infection. J Biomed Biotechnol 2010, 159152. 10.1155/2010/159152.

17. Laidlaw, B.J., Zhang, N., Marshall, H.D., Staron, M.M., Guan, T., Hu, Y., Cauley, L.S., Craft, J., and Kaech, S.M. (2014). CD4+ T cell help guides formation of CD103+ lung-resident memory CD8+ T cells during influenza viral infection. Immunity 41, 633–645. 10.1016/j.immuni.2014.09.007.

18. Cullen, J.G., McQuilten, H.A., Quinn, K.M., Olshansky, M., Russ, B.E., Morey, A., Wei, S., Prier, J.E., La Gruta, N.L., Doherty, P.C., and Turner, S.J. (2019). CD4(+) T help promotes influenza virus-specific CD8(+) T cell memory by limiting metabolic dysfunction. Proc Natl Acad Sci U S A 116, 4481–4488. 10.1073/pnas.1808849116.

19. Northrop, J.K., Wells, A.D., and Shen, H. (2008). Cutting edge: chromatin remodeling as a molecular basis for the enhanced functionality of memory CD8 T cells. J Immunol 181, 865–868.

20. Northrop, J.K., Thomas, R.M., Wells, A.D., and Shen, H. (2006). Epigenetic remodeling of the IL-2 and IFN-gamma loci in memory CD8 T cells is influenced by CD4 T cells. J Immunol 177, 1062–1069. 10.4049/jimmunol.177.2.1062.

21. Provine, N.M., Larocca, R.A., Aid, M., Penaloza-MacMaster, P., Badamchi-Zadeh, A., Borducchi, E.N., Yates, K.B., Abbink, P., Kirilova, M., Ng’ang’a, D., et al. (2016). Immediate Dysfunction of Vaccine-Elicited CD8+ T Cells Primed in the Absence of CD4+ T Cells. J Immunol 197, 1809–1822. 10.4049/jimmunol.1600591.

22. Busselaar, J., Tian, S., van Eenennaam, H., and Borst, J. (2020). Helpless Priming Sends CD8(+) T Cells on the Road to Exhaustion. Frontiers in immunology 11, 592569. 10.3389/fimmu.2020.592569.

23. Lu, Y.J., Barreira-Silva, P., Boyce, S., Powers, J., Cavallo, K., and Behar, S.M. (2021). CD4 T cell help prevents CD8 T cell exhaustion and promotes control of Mycobacterium tuberculosis infection. Cell Rep 36, 109696. 10.1016/j.celrep.2021.109696.

24. Roberts, A.D., Ely, K.H., and Woodland, D.L. (2005). Differential contributions of central and effector memory T cells to recall responses. J Exp Med 202, 123–133.

25. Hikono, H., Kohlmeier, J.E., Takamura, S., Wittmer, S.T., Roberts, A.D., and Woodland, D.L. (2007). Activation phenotype, rather than central- or effector-memory phenotype, predicts the recall efficacy of memory CD8+ T cells. J Exp Med 204, 1625–1636.

26. Kaech, S.M., and Wherry, E.J. (2007). Heterogeneity and cell-fate decisions in effector and memory CD8+ T cell differentiation during viral infection. Immunity 27, 393–405.

27. Martin, M.D., Kim, M.T., Shan, Q., Sompallae, R., Xue, H.H., Harty, J.T., and Badovinac, V.P. (2015). Phenotypic and Functional Alterations in Circulating Memory CD8 T Cells with Time after Primary Infection. PLoS Pathog 11, e1005219. 10.1371/journal.ppat.1005219.

28. Eberlein, J., Davenport, B., Nguyen, T., Victorino, F., Haist, K., Jhun, K., Karimpour-Fard, A., Hunter, L., Kedl, R., Clambey, E.T., and Homann, D. (2016). Aging promotes acquisition of naive-like CD8+ memory T cell traits and enhanced functionalities. J Clin Invest 126, 3942–3960. 10.1172/JCI88546.

29. Davenport, B., Eberlein, J., van der Heide, V., Jhun, K., Nguyen, T.T., Victorino, F., Trotta, A., Chipuk, J., Yi, Z., Zhang, W., et al. (2019). Aging of Antiviral CD8(+) Memory T Cells Fosters Increased Survival, Metabolic Adaptations, and Lymphoid Tissue Homing. J Immunol 202, 460–475. 10.4049/jimmunol.1801277.

30. Davenport, B., Eberlein, J., Nguyen, T.T., Victorino, F., Jhun, K., Abuirqeba, H., van der Heide, V., Heeger, P., and Homann, D. (2019). Aging boosts antiviral CD8+T cell memory through improved engagement of diversified recall response determinants. PLoS Pathog 15, e1008144. 10.1371/journal.ppat.1008144.

31. Davenport, B., Eberlein, J., Nguyen, T.T., Victorino, F., van der Heide, V., Kuleshov, M., Ma’ayan, A., Kedl, R., and Homann, D. (2020). Chemokine Signatures of Pathogen-Specific T Cells II: Memory T Cells in Acute and Chronic Infection. J Immunol 205, 2188–2206. 10.4049/jimmunol.2000254.

32. Gerlach, C., Moseman, E.A., Loughhead, S.M., Alvarez, D., Zwijnenburg, A.J., Waanders, L., Garg, R., de la Torre, J.C., and von Andrian, U.H. (2016). The Chemokine Receptor CX3CR1 Defines Three Antigen-Experienced CD8 T Cell Subsets with Distinct Roles in Immune Surveillance and Homeostasis. Immunity 45, 1270–1284. 10.1016/j.immuni.2016.10.018.

33. Zwijnenburg, A.J., Pokharel, J., Varnaite, R., Zheng, W., Hoffer, E., Shryki, I., Comet, N.R., Ehrstrom, M., Gredmark-Russ, S., Eidsmo, L., and Gerlach, C. (2023). Graded expression of the chemokine receptor CX3CR1 marks differentiation states of human and murine T cells and enables cross-species interpretation. Immunity 56, 1955–1974 e1910. 10.1016/j.immuni.2023.06.025.

34. Gill, A.L., Hudson, W.H., Valanparambil, R.M., Ahn, E., McGuire, D.J., Wieland, A., McManus, D.T., Kissick, H.T., Akondy, R.S., and Ahmed, R. (2023). Longitudinal Analysis of the Phenotype, Transcriptional Profile, and Anatomic Location of Memory CD8 T Cell Subsets after Acute Viral Infection. J Virol 97, e0155622. 10.1128/jvi.01556-22.

35. O’Sullivan, D. (2019). The metabolic spectrum of memory T cells. Immunol Cell Biol 97, 636–646. 10.1111/imcb.12274.

36. Miller, J.D., van der Most, R.G., Akondy, R.S., Glidewell, J.T., Albott, S., Masopust, D., Murali-Krishna, K., Mahar, P.L., Edupuganti, S., Lalor, S., et al. (2008). Human effector and memory CD8+ T cell responses to smallpox and yellow fever vaccines. Immunity 28, 710–722. S1074-7613(08)00194-5 [pii] 10.1016/j.immuni.2008.02.020.

37. Akondy, R.S., Monson, N.D., Miller, J.D., Edupuganti, S., Teuwen, D., Wu, H., Quyyumi, F., Garg, S., Altman, J.D., Del Rio, C., et al. (2009). The yellow fever virus vaccine induces a broad and polyfunctional human memory CD8+ T cell response. J Immunol 183, 7919–7930. 10.4049/jimmunol.0803903.

38. Akondy, R.S., Fitch, M., Edupuganti, S., Yang, S., Kissick, H.T., Li, K.W., Youngblood, B.A., Abdelsamed, H.A., McGuire, D.J., Cohen, K.W., et al. (2017). Origin and differentiation of human memory CD8 T cells after vaccination. Nature 552, 362–367. 10.1038/nature24633.

39. Youngblood, B., Hale, J.S., Kissick, H.T., Ahn, E., Xu, X., Wieland, A., Araki, K., West, E.E., Ghoneim, H.E., Fan, Y., et al. (2017). Effector CD8 T cells dedifferentiate into long-lived memory cells. Nature 552, 404–409. 10.1038/nature25144.

40. Ahmed, R., Butler, L.D., and Bhatti, L. (1988). T4+ T helper cell function in vivo: differential requirement for induction of antiviral cytotoxic T-cell and antibody responses. J Virol 62, 2102–2106. 10.1128/JVI.62.6.2102-2106.1988.

41. Rahemtulla, A., Fung-Leung, W.P., Schilham, M.W., Kundig, T.M., Sambhara, S.R., Narendran, A., Arabian, A., Wakeham, A., Paige, C.J., Zinkernagel, R.M., and, et al. (1991). Normal development and function of CD8+ cells but markedly decreased helper cell activity in mice lacking CD4. Nature 353, 180–184. 10.1038/353180a0.

42. Matloubian, M., Concepcion, R.J., and Ahmed, R. (1994). CD4+ T cells are required to sustain CD8+ cytotoxic T-cell responses during chronic viral infection. J Virol 68, 8056–8063. 10.1128/JVI.68.12.8056-8063.1994.

43. Tishon, A., Lewicki, H., Rall, G., Von Herrath, M., and Oldstone, M.B. (1995). An essential role for type 1 interferon-gamma in terminating persistent viral infection. Virology 212, 244–250.

44. von Herrath, M.G., Yokoyama, M., Dockter, J., Oldstone, M.B., and Whitton, J.L. (1996). CD4-deficient mice have reduced levels of memory cytotoxic T lymphocytes after immunization and show diminished resistance to subsequent virus challenge. J Virol 70, 1072–1079.

45. Badovinac, V.P., Haring, J.S., and Harty, J.T. (2007). Initial T cell receptor transgenic cell precursor frequency dictates critical aspects of the CD8(+) T cell response to infection. Immunity 26, 827–841.

46. Olson, J.A., McDonald-Hyman, C., Jameson, S.C., and Hamilton, S.E. (2013). Effector-like CD8(+) T cells in the memory population mediate potent protective immunity. Immunity 38, 1250–1260. 10.1016/j.immuni.2013.05.009.

47. Herndler-Brandstetter, D., Ishigame, H., Shinnakasu, R., Plajer, V., Stecher, C., Zhao, J., Lietzenmayer, M., Kroehling, L., Takumi, A., Kometani, K., et al. (2018). KLRG1(+) Effector CD8(+) T Cells Lose KLRG1, Differentiate into All Memory T Cell Lineages, and Convey Enhanced Protective Immunity. Immunity 48, 716–729 e718. 10.1016/j.immuni.2018.03.015.

48. Milner, J.J., Nguyen, H., Omilusik, K., Reina-Campos, M., Tsai, M., Toma, C., Delpoux, A., Boland, B.S., Hedrick, S.M., Chang, J.T., and Goldrath, A.W. (2020). Delineation of a molecularly distinct terminally differentiated memory CD8 T cell population. Proc Natl Acad Sci U S A 117, 25667–25678. 10.1073/pnas.2008571117.

49. Evrard, M., Becht, E., Fonseca, R., Obers, A., Park, S.L., Ghabdan-Zanluqui, N., Schroeder, J., Christo, S.N., Schienstock, D., Lai, J., et al. (2023). Single-cell protein expression profiling resolves circulating and resident memory T cell diversity across tissues and infection contexts. Immunity 56, 1664–1680 e1669. 10.1016/j.immuni.2023.06.005.

50. Kohlmeier, J.E., Reiley, W.W., Perona-Wright, G., Freeman, M.L., Yager, E.J., Connor, L.M., Brincks, E.L., Cookenham, T., Roberts, A.D., Burkum, C.E., et al. (2011). Inflammatory chemokine receptors regulate CD8(+) T cell contraction and memory generation following infection. J Exp Med 208, 1621–1634. jem.20102110 [pii] 10.1084/jem.20102110.

51. Slutter, B., Pewe, L.L., Kaech, S.M., and Harty, J.T. (2013). Lung airway-surveilling CXCR3(hi) memory CD8(+) T cells are critical for protection against influenza A virus. Immunity 39, 939–948. 10.1016/j.immuni.2013.09.013.

52. Sallusto, F., Lenig, D., Forster, R., Lipp, M., and Lanzavecchia, A. (1999). Two subsets of memory T lymphocytes with distinct homing potentials and effector functions. Nature 401, 708–712.

53. Renkema, K.R., Huggins, M.A., Borges da Silva, H., Knutson, T.P., Henzler, C.M., and Hamilton, S.E. (2020). KLRG1(+) Memory CD8 T Cells Combine Properties of Short-Lived Effectors and Long-Lived Memory. J Immunol 205, 1059–1069. 10.4049/jimmunol.1901512.

54. Wherry, E.J., Teichgraber, V., Becker, T.C., Masopust, D., Kaech, S.M., Antia, R., von Andrian, U.H., and Ahmed, R. (2003). Lineage relationship and protective immunity of memory CD8 T cell subsets. Nat Immunol 4, 225–234.

55. Bachmann, M.F., Wolint, P., Schwarz, K., Jager, P., and Oxenius, A. (2005). Functional properties and lineage relationship of CD8+ T cell subsets identified by expression of IL-7 receptor alpha and CD62L. J Immunol 175, 4686–4696. 10.4049/jimmunol.175.7.4686.

56. Joshi, N.S., Cui, W., Chandele, A., Lee, H.K., Urso, D.R., Hagman, J., Gapin, L., and Kaech, S.M. (2007). Inflammation Directs Memory Precursor and Short-Lived Effector CD8(+) T Cell Fates via the Graded Expression of T-bet Transcription Factor. Immunity 27, 281–295.

57. Utzschneider, D.T., Delpoux, A., Wieland, D., Huang, X., Lai, C.Y., Hofmann, M., Thimme, R., and Hedrick, S.M. (2018). Active Maintenance of T Cell Memory in Acute and Chronic Viral Infection Depends on Continuous Expression of FOXO1. Cell Rep 22, 3454–3467. 10.1016/j.celrep.2018.03.020.

58. Zhou, X., Yu, S., Zhao, D.M., Harty, J.T., Badovinac, V.P., and Xue, H.H. (2010). Differentiation and persistence of memory CD8(+) T cells depend on T cell factor 1. Immunity 33, 229–240. 10.1016/j.immuni.2010.08.002.

59. McLane, L.M., Abdel-Hakeem, M.S., and Wherry, E.J. (2019). CD8 T Cell Exhaustion During Chronic Viral Infection and Cancer. Annu Rev Immunol. 10.1146/annurev-immunol-041015-055318.

60. Marzo, A.L., Klonowski, K.D., Le Bon, A., Borrow, P., Tough, D.F., and Lefrancois, L. (2005). Initial T cell frequency dictates memory CD8+ T cell lineage commitment. Nat Immunol 6, 793–799.

61. Blattman, J.N., Antia, R., Sourdive, D.J., Wang, X., Kaech, S.M., Murali-Krishna, K., Altman, J.D., and Ahmed, R. (2002). Estimating the precursor frequency of naive antigen-specific CD8 T cells. J Exp Med 195, 657–664.

62. Kaech, S.M., Tan, J.T., Wherry, E.J., Konieczny, B.T., Surh, C.D., and Ahmed, R. (2003). Selective expression of the interleukin 7 receptor identifies effector CD8 T cells that give rise to long-lived memory cells. Nat Immunol 4, 1191–1198.

63. Sarkar, S., Kalia, V., Haining, W.N., Konieczny, B.T., Subramaniam, S., and Ahmed, R. (2008). Functional and genomic profiling of effector CD8 T cell subsets with distinct memory fates. J Exp Med 205, 625–640. jem.20071641 [pii] 10.1084/jem.20071641.

64. Plumlee, C.R., Sheridan, B.S., Cicek, B.B., and Lefrancois, L. (2013). Environmental cues dictate the fate of individual CD8+ T cells responding to infection. Immunity 39, 347–356. 10.1016/j.immuni.2013.07.014.

65. Ahrends, T., Spanjaard, A., Pilzecker, B., Babala, N., Bovens, A., Xiao, Y., Jacobs, H., and Borst, J. (2017). CD4(+) T Cell Help Confers a Cytotoxic T Cell Effector Program Including Coinhibitory Receptor Downregulation and Increased Tissue Invasiveness. Immunity 47, 848–861 e845. 10.1016/j.immuni.2017.10.009.

66. Becht, E., McInnes, L., Healy, J., Dutertre, C.A., Kwok, I.W.H., Ng, L.G., Ginhoux, F., and Newell, E.W. (2018). Dimensionality reduction for visualizing single-cell data using UMAP. Nat Biotechnol. 10.1038/nbt.4314.

67. Traag, V.A., Waltman, L., and van Eck, N.J. (2019). From Louvain to Leiden: guaranteeing well-connected communities. Sci Rep 9, 5233. 10.1038/s41598-019-41695-z.

68. Giles, J.R., Ngiow, S.F., Manne, S., Baxter, A.E., Khan, O., Wang, P., Staupe, R., Abdel-Hakeem, M.S., Huang, H., Mathew, D., et al. (2022). Shared and distinct biological circuits in effector, memory and exhausted CD8(+) T cells revealed by temporal single-cell transcriptomics and epigenetics. Nat Immunol 23, 1600–1613. 10.1038/s41590-022-01338-4.

69. Squair, J.W., Gautier, M., Kathe, C., Anderson, M.A., James, N.D., Hutson, T.H., Hudelle, R., Qaiser, T., Matson, K.J.E., Barraud, Q., et al. (2021). Confronting false discoveries in single-cell differential expression. Nature communications 12, 5692. 10.1038/s41467-021-25960-2.

70. Murphy, A.E., and Skene, N.G. (2022). A balanced measure shows superior performance of pseudobulk methods in single-cell RNA-sequencing analysis. Nature communications 13, 7851. 10.1038/s41467-022-35519-4.

71. Danilo, M., Chennupati, V., Silva, J.G., Siegert, S., and Held, W. (2018). Suppression of Tcf1 by Inflammatory Cytokines Facilitates Effector CD8 T Cell Differentiation. Cell Rep 22, 2107–2117. 10.1016/j.celrep.2018.01.072.

72. Pais Ferreira, D., Silva, J.G., Wyss, T., Fuertes Marraco, S.A., Scarpellino, L., Charmoy, M., Maas, R., Siddiqui, I., Tang, L., Joyce, J.A., et al. (2020). Central memory CD8(+) T cells derive from stem-like Tcf7(hi) effector cells in the absence of cytotoxic differentiation. Immunity 53, 985–1000 e1011. 10.1016/j.immuni.2020.09.005.

73. Silva, J.G., Pais Ferreira, D., Dumez, A., Wyss, T., Veber, R., Danilo, M., Pinschewer, D.D., Charmoy, M., and Held, W. (2023). Emergence and fate of stem cell-like Tcf7(+) CD8(+) T cells during a primary immune response to viral infection. Sci Immunol 8, eadh3113. 10.1126/sciimmunol.adh3113.

74. Johnnidis, J.B., Muroyama, Y., Ngiow, S.F., Chen, Z., Manne, S., Cai, Z., Song, S., Platt, J.M., Schenkel, J.M., Abdel-Hakeem, M., et al. (2021). Inhibitory signaling sustains a distinct early memory CD8(+) T cell precursor that is resistant to DNA damage. Sci Immunol 6. 10.1126/sciimmunol.abe3702.

75. Harrington, L.E., Galvan, M., Baum, L.G., Altman, J.D., and Ahmed, R. (2000). Differentiating between memory and effector CD8 T cells by altered expression of cell surface O-glycans. J Exp Med 191, 1241–1246.

76. Yang, Q., Li, F., Harly, C., Xing, S., Ye, L., Xia, X., Wang, H., Wang, X., Yu, S., Zhou, X., et al. (2015). TCF-1 upregulation identifies early innate lymphoid progenitors in the bone marrow. Nat Immunol 16, 1044–1050. 10.1038/ni.3248.

77. Schweier, O., Aichele, U., Marx, A.F., Straub, T., Verbeek, J.S., Pinschewer, D.D., and Pircher, H. (2019). Residual LCMV antigen in transiently CD4(+) T cell-depleted mice induces high levels of virus-specific antibodies but only limited B-cell memory. Eur J Immunol 49, 626–637. 10.1002/eji.201847772.

78. Bachmann, M.F., Hunziker, L., Zinkernagel, R.M., Storni, T., and Kopf, M. (2004). Maintenance of memory CTL responses by T helper cells and CD40-CD40 ligand: antibodies provide the key. Eur J Immunol 34, 317–326. 10.1002/eji.200324717.

79. Whitmire, J.K., Eam, B., and Whitton, J.L. (2008). Tentative T cells: memory cells are quick to respond, but slow to divide. PLoS Pathog 4, e1000041.

80. Mehlhop-Williams, E.R., and Bevan, M.J. (2014). Memory CD8+ T cells exhibit increased antigen threshold requirements for recall proliferation. J Exp Med 211, 345–356. 10.1084/jem.20131271.

81. Snell, L.M., MacLeod, B.L., Law, J.C., Osokine, I., Elsaesser, H.J., Hezaveh, K., Dickson, R.J., Gavin, M.A., Guidos, C.J., McGaha, T.L., and Brooks, D.G. (2018). CD8(+) T Cell Priming in Established Chronic Viral Infection Preferentially Directs Differentiation of Memory-like Cells for Sustained Immunity. Immunity 49, 678–694 e675. 10.1016/j.immuni.2018.08.002.

82. Sun, J.C., Williams, M.A., and Bevan, M.J. (2004). CD4+ T cells are required for the maintenance, not programming, of memory CD8+ T cells after acute infection. Nat Immunol 5, 927–933.

83. Choo, D.K., Murali-Krishna, K., Anita, R., and Ahmed, R. (2010). Homeostatic turnover of virus-specific memory CD8 T cells occurs stochastically and is independent of CD4 T cell help. J Immunol 185, 3436–3444. 10.4049/jimmunol.1001421.

84. Puglielli, M.T., Zajac, A.J., van der Most, R.G., Dzuris, J.L., Sette, A., Altman, J.D., and Ahmed, R. (2001). In vivo selection of a lymphocytic choriomeningitis virus variant that affects recognition of the GP33-43 epitope by H-2Db but not H-2Kb. J Virol 75, 5099–5107. 10.1128/JVI.75.11.5099-5107.2001.

85. Lan, X., Zebley, C.C., and Youngblood, B. (2023). Cellular and molecular waypoints along the path of T cell exhaustion. Sci Immunol 8, eadg3868. 10.1126/sciimmunol.adg3868.

86. Klenerman, P. (2018). The (gradual) rise of memory inflation. Immunol Rev 283, 99–112. 10.1111/imr.12653.

87. Kim, T.S., Sun, J., and Braciale, T.J. (2011). T cell responses during influenza infection: getting and keeping control. Trends Immunol 32, 225–231. 10.1016/j.it.2011.02.006.

88. Proal, A.D., VanElzakker, M.B., Aleman, S., Bach, K., Boribong, B.P., Buggert, M., Cherry, S., Chertow, D.S., Davies, H.E., Dupont, C.L., et al. (2023). SARS-CoV-2 reservoir in post-acute sequelae of COVID-19 (PASC). Nat Immunol 24, 1616–1627. 10.1038/s41590-023-01601-2.

89. Yin, K., Peluso, M.J., Luo, X., Thomas, R., Shin, M.G., Neidleman, J., Andrew, A., Young, K.C., Ma, T., Hoh, R., et al. (2024). Long COVID manifests with T cell dysregulation, inflammation and an uncoordinated adaptive immune response to SARS-CoV-2. Nat Immunol. 10.1038/s41590-023-01724-6.

90. Abdel-Hakeem, M.S., Manne, S., Beltra, J.C., Stelekati, E., Chen, Z., Nzingha, K., Ali, M.A., Johnson, J.L., Giles, J.R., Mathew, D., et al. (2021). Epigenetic scarring of exhausted T cells hinders memory differentiation upon eliminating chronic antigenic stimulation. Nat Immunol 22, 1008–1019. 10.1038/s41590-021-00975-5.

91. Ahrends, T., Busselaar, J., Severson, T.M., Babala, N., de Vries, E., Bovens, A., Wessels, L., van Leeuwen, F., and Borst, J. (2019). CD4(+) T cell help creates memory CD8(+) T cells with innate and help-independent recall capacities. Nature communications 10, 5531. 10.1038/s41467-019-13438-1.

92. van der Heide, V., Humblin, E., Vaidya, A., and Kamphorst, A.O. (2022). Advancing beyond the twists and turns of T cell exhaustion in cancer. Sci Transl Med 14, eabo4997. 10.1126/scitranslmed.abo4997.

93. Laidlaw, B.J., Cui, W., Amezquita, R.A., Gray, S.M., Guan, T., Lu, Y., Kobayashi, Y., Flavell, R.A., Kleinstein, S.H., Craft, J., and Kaech, S.M. (2015). Production of IL-10 by CD4(+) regulatory T cells during the resolution of infection promotes the maturation of memory CD8(+) T cells. Nat Immunol 16, 871–879. 10.1038/ni.3224.

94. Kalia, V., Penny, L.A., Yuzefpolskiy, Y., Baumann, F.M., and Sarkar, S. (2015). Quiescence of Memory CD8(+) T Cells Is Mediated by Regulatory T Cells through Inhibitory Receptor CTLA-4. Immunity 42, 1116–1129. 10.1016/j.immuni.2015.05.023.

95. Recher, M., Lang, K.S., Navarini, A., Hunziker, L., Lang, P.A., Fink, K., Freigang, S., Georgiev, P., Hangartner, L., Zellweger, R., et al. (2007). Extralymphatic virus sanctuaries as a consequence of potent T-cell activation. Nat Med 13, 1316–1323. 10.1038/nm1670.

96. Gullicksrud, J.A., Li, F., Xing, S., Zeng, Z., Peng, W., Badovinac, V.P., Harty, J.T., and Xue, H.H. (2017). Differential Requirements for Tcf1 Long Isoforms in CD8(+) and CD4(+) T Cell Responses to Acute Viral Infection. J Immunol 199, 911–919. 10.4049/jimmunol.1700595.

97. Ahmed, R., Salmi, A., Butler, L.D., Chiller, J.M., and Oldstone, M.B. (1984). Selection of genetic variants of lymphocytic choriomeningitis virus in spleens of persistently infected mice. Role in suppression of cytotoxic T lymphocyte response and viral persistence. J Exp Med 160, 521–540.

98. Ahmed, R., and Oldstone, M.B. (1988). Organ-specific selection of viral variants during chronic infection. J Exp Med 167, 1719–1724. 10.1084/jem.167.5.1719.

99. Sanchez, A.B., and de la Torre, J.C. (2006). Rescue of the prototypic Arenavirus LCMV entirely from plasmid. Virology 350, 370–380. 10.1016/j.virol.2006.01.012.

100. Emonet, S.F., Garidou, L., McGavern, D.B., and de la Torre, J.C. (2009). Generation of recombinant lymphocytic choriomeningitis viruses with trisegmented genomes stably expressing two additional genes of interest. Proc Natl Acad Sci U S A 106, 3473–3478. 10.1073/pnas.0900088106.

101. Iwasaki, M., Ngo, N., Cubitt, B., Teijaro, J.R., and de la Torre, J.C. (2015). General Molecular Strategy for Development of Arenavirus Live-Attenuated Vaccines. J Virol 89, 12166–12177. 10.1128/JVI.02075-15.

102. Heinen, A.P., Wanke, F., Moos, S., Attig, S., Luche, H., Pal, P.P., Budisa, N., Fehling, H.J., Waisman, A., and Kurschus, F.C. (2014). Improved method to retain cytosolic reporter protein fluorescence while staining for nuclear proteins. Cytometry A 85, 621–627. 10.1002/cyto.a.22451.

103. Roederer, M. (2011). Interpretation of cellular proliferation data: avoid the panglossian. Cytometry A 79, 95–101. 10.1002/cyto.a.21010.

104. Jabbari, A., and Harty, J.T. (2006). Simultaneous assessment of antigen-stimulated cytokine production and memory subset composition of memory CD8 T cells. J Immunol Methods 313, 161–168. S0022-1759(06)00113-X [pii] 10.1016/j.jim.2006.04.005.

105. Zheng, G.X., Terry, J.M., Belgrader, P., Ryvkin, P., Bent, Z.W., Wilson, R., Ziraldo, S.B., Wheeler, T.D., McDermott, G.P., Zhu, J., et al. (2017). Massively parallel digital transcriptional profiling of single cells. Nature communications 8, 14049. 10.1038/ncomms14049.

106. Wolf, F.A., Angerer, P., and Theis, F.J. (2018). SCANPY: large-scale single-cell gene expression data analysis. Genome Biol 19, 15. 10.1186/s13059-017-1382-0.

107. Aran, D., Looney, A.P., Liu, L., Wu, E., Fong, V., Hsu, A., Chak, S., Naikawadi, R.P., Wolters, P.J., Abate, A.R., et al. (2019). Reference-based analysis of lung single-cell sequencing reveals a transitional profibrotic macrophage. Nat Immunol 20, 163–172. 10.1038/s41590-018-0276-y.

108. Badia, I.M.P., Velez Santiago, J., Braunger, J., Geiss, C., Dimitrov, D., Muller-Dott, S., Taus, P., Dugourd, A., Holland, C.H., Ramirez Flores, R.O., and Saez-Rodriguez, J. (2022). decoupleR: ensemble of computational methods to infer biological activities from omics data. Bioinform Adv 2, vbac016. 10.1093/bioadv/vbac016.

