## Supplemental information for "Functional impairment of “helpless” CD8^+^ memory T cells is transient and driven by prolonged but finite cognate antigen presentation"

#### Figure S1 (related to Figure 1)

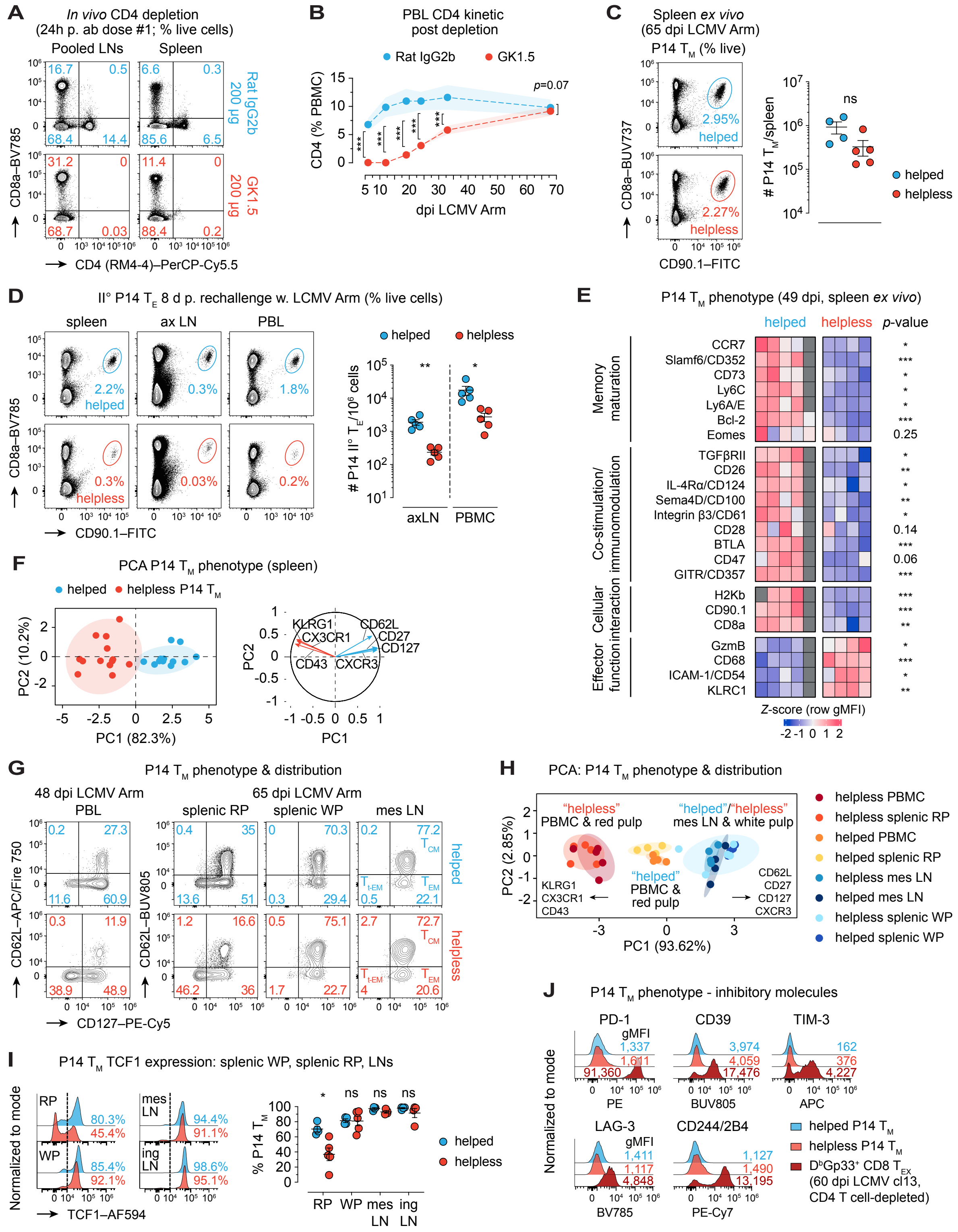

***Figure S1*** **(related to Figure 1)*.* Helpless P14 $T_M$ are broadly immature**

(**A**-**B**) Efficacy and duration of transient $CD4^+$ T cell depletion. Shown are (**A**) representative flow cytometry plots of percentage $CD4^+$ T among live cells in naïve C57BL/6 (B6) spleens as well as pooled inguinal (ing)/mesenteric (mes)/brachial lymph nodes (LNs) after one out of two doses of anti-CD4 antibody (clone GK1.5) *vs*. isotype control (Rat IgG2b), and (**B**) time course indicating percentages of peripheral blood (PBL) $CD4^+$ T cells after transient depletion until full restoration in helpless *vs*. helped LCMV-immune P14 chimeras. (**C**) (Left) representative flow cytometry plots and (right) quantification of splenic P14 $T_M$ at 65 dpi *ex vivo*. (**D**) II° P14 $T_E$ responses by P14 $T_M$ 8d post rechallenge (RC) with LCMV Arm by (left) representative flow cytometry plots of spleens, axillary lymph nodes (ax LN) as well as PBL, and (right) enumeration (see ***Figures 1A*** and ***1B***). (**E**) Comparative heatmap visualization and statistics of *Z*-score normalized row geometric mean fluorescence intensity (gMFI) values of splenic P14 $T_M$-expressed molecules associated with $T_M$ maturation and function at 49 dpi. Missing values are indicated by dark grey color. (**F**) Principal component analysis (PCA) clustering (left) of helped and helpless P14 $T_M$ phenotypes based on markers in corresponding loading plot (right). (**G**) Representative flow cytometry plots showing frequencies of $CD127^+CD62L^+$ central ($T_{CM}$), $CD127^+CD62L^-$ transitional effector ($T_{EM}$), and $CD127^-CD62L^-$ terminally differentiated effector memory ($T_{t\text{-}EM}$) subsets as recently described[1] among P14 $T_M$ in peripheral blood (PBL), splenic red (RP) and white pulp (WP), as well as mes LN at 48 or 65 dpi, respectively. (**H**) PCA based on percentual expression of P14 $T_M$ markers used in (**F**) comparing helped and helpless P14 $T_M$ phenotypes across peripheral blood mononuclear cells (PBMC), splenic RP, mesLN, and splenic WP. (**I**) (Left) representative histogram overlays and (right) quantification of splenic P14 $T_M$-expressed TCF1 in RP, WP, mes LN, and ing LN. (**J**) Representative histogram overlays comparing expression by gMFI of "traditional" exhaustion markers on helped (blue) and helpless (red) splenic P14 $T_M$ at 60 dpi LCMV Arm *vs*. *bona fide* exhausted (dark red) $D^bGp_{33}{}^+$ LCMV-specific $CD8^+$ T cells ($CD8^+$ $T_{EX}$) 60d after chronic LCMV cl13 infection of $CD4^+$ T cell-depleted B6 mice. Graphs from one representative out of at least two independent experiments plot individual helped (blue) and helpless (red) mice with n=9-10 animals/group/time point and mean±95% confidence intervals in (**B**) as well as n=4-5 animals/group/time point/experiment and mean±SEM in (**C**), (**D**), and (**I**). Heatmap data in (**E**) show individual mice representative of one-two experimens with n=3-4mice/group/experiment; Bcl-2 and Eomes data from different experiment representative of at least 2 independent experiments with n=4-5 mice/group/experiment. PCA with 95% confidence ellipses in (**F**) are pooled from three independent experiments with n=4-5 mice/group/experiment. PCA with 80% confidence ellipses in (**H**) plots individual mice from one representative experiment. Histogram overlays in (**J**) are representative of n=4-5 mice/group from one representative experiment.

*$p<0.05$, **$p<0.01$, and ***$p<0.001$ by unpaired Student's *t* test or Mann Whitney U test. ns, non-significant.

#### Figure S2 (related to Figures 2 & 3)

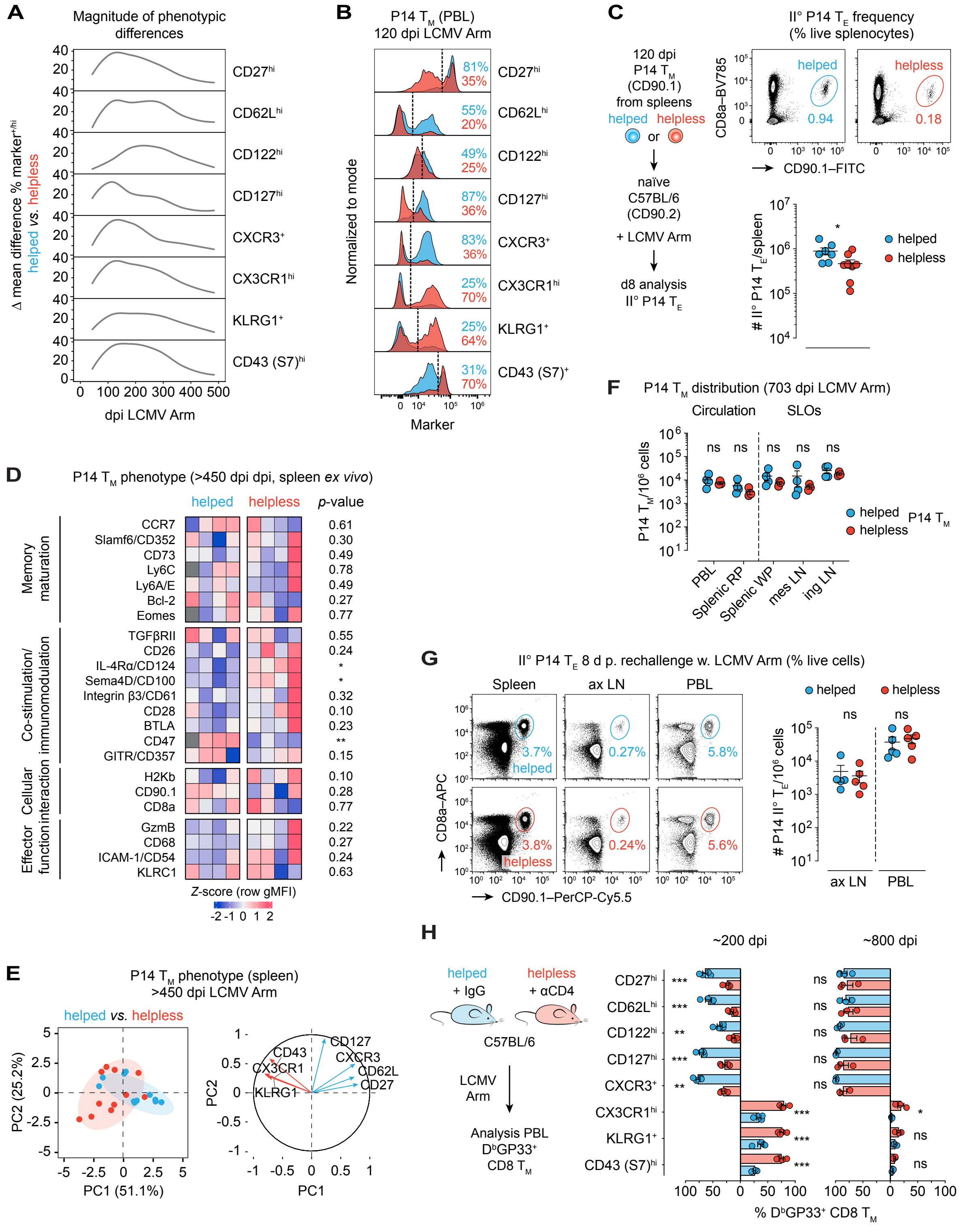

***Figure S2*** **(related to Figures 2 & 3)*. Prolonged but complete $T_M$ maturation in helpless mice***

(**A**) Time course analysis depicting magnitude of differences in mean expression levels of CD27, CD62L, CD122, CD127, CXCR3, CX3CR1, and CD43 (clone S7) by peripheral blood helped *vs*. helpless P14 $T_M$ spanning ~d49-500 after LCMV Arm. (**B**) Representative histogram overlays of peripheral blood P14 $T_M$ phenotypes at ~120 dpi. (**C**) (Top) representative flow cytometry plots and (bottom) enumeration of recall response to LCMV Arm by splenic P14 $T_M$ at ~120 dpi. (**D**) Comparative heatmap visualization and statistics of *Z*-score normalized row gMFI values of splenic P14 $T_M$-expressed molecules associated with $T_M$ maturation and function at >450 dpi (see ***Figure S1E***). Missing values are indicated by dark grey color. (**E**) (Left) PCA of helped and helpless P14 $T_M$ phenotypes in spleens > 450 dpi (see ***Figure S1F***).(**F**) Quantification of P14 $T_M$ numbers in circulation (PBL, RP) and selected SLOs (WP, mes LN, ing LN) (see ***Figure 3E***). (**G**) II° P14 $T_E$ responses by >450 dpi splenic P14 $T_M$ 8d post rechallenge (RC) with LCMV Arm by (left) representative flow cytometry plots of spleens, ax LNs, and PBL as well as (right) enumeration (see ***Figure 3J***). (**H**) C57BL/6 mice were treated with isotype (IgG/helped) or anti-$CD4^+$ T cell depleting antibody (αCD4/helpless) before LCMV Arm infection. (Left) phenotypic expression profiles of PBL $D^bGP_{33}^+$ $CD8^+$ $T_M$ at ~200 (left) and at ~800 dpi (right). Graphs from one representative out of two independent experiments plot individual helped (blue) and helpless (red) mice with n=7-8 animals/group/time point and mean±SEM in (**C**), n=3-5 animals/group/time point/experiment and mean±SEM in (**F**), (**G**), and (**H**). For time course data in (**A**), ∆ mean difference % $marker^{+/hi}$ was calculated by subtracting lower from higher mean values for each marker with n=5-14 mice/group/time point. Heatmap data in (**D**) are individual mice from one representative experiment. PCA with 95% confidence ellipses in (**E**) are pooled from three independent experiments with n=3-4 mice/group/experiment. *$p$<0.05, **$p$<0.01, and ***$p$<0.001 by unpaired Student's *t* test or Mann Whitney U test. ns, non-significant.

#### Figure S3 (related to Figure 4)

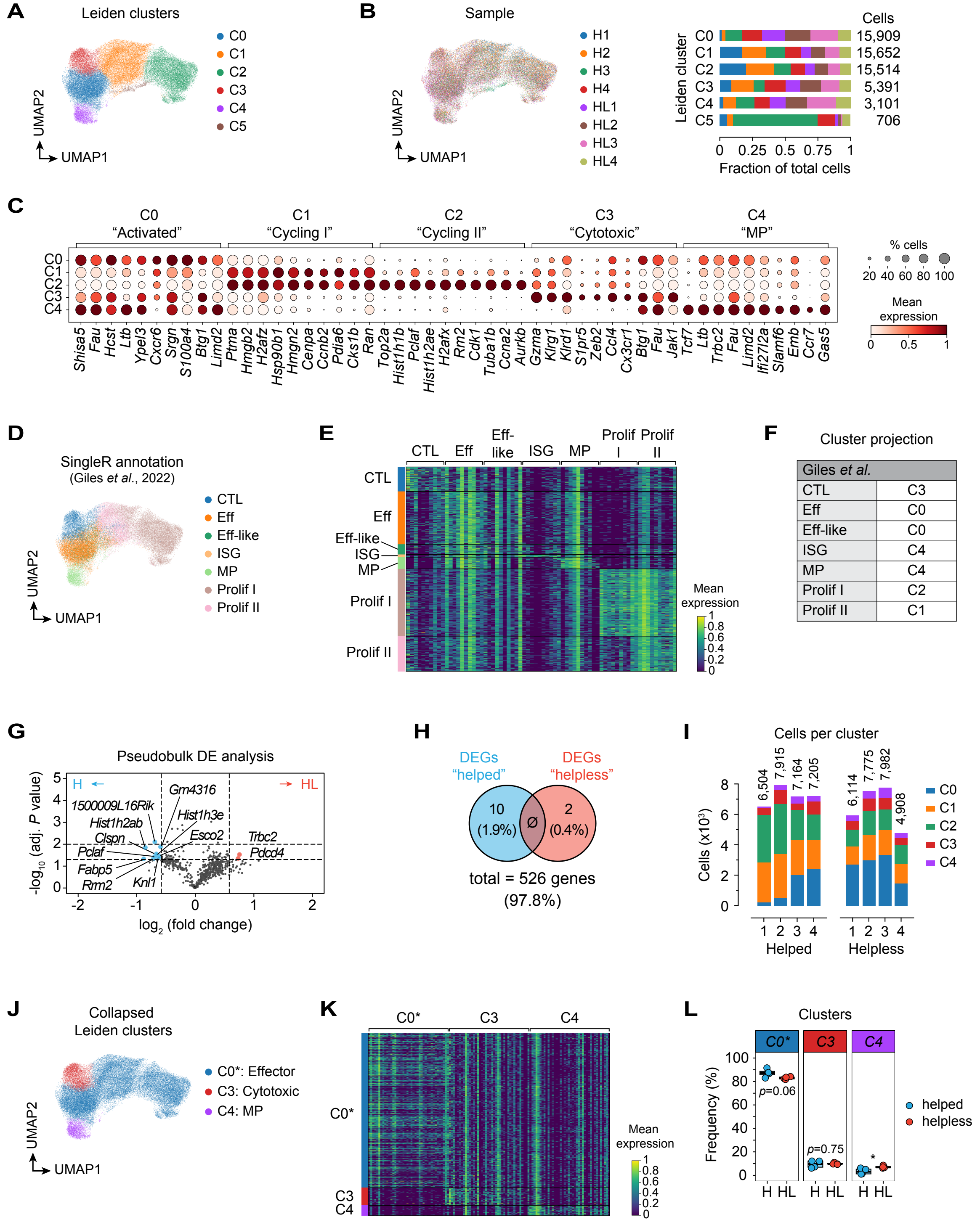

***Figure S3*** **(related to Figure 4)*.* Minor transcriptomic differences between helped and helpless P14 $T_E$**

(**A**-**L**) Helped and helpless splenic P14 $T_E$ were sort-purifed at 7 dpi with LCMV Arm and analyzed by scRNA-seq (n=4/group). (**A**-**B**) (**A**) UMAP projection with six Leiden clusters including sample-biased cluster C5, and (**B**) with distribution of individual helped and helpless samples (left) as well as relative fraction and total number of cells per cluster (right). Note that sample-biased cluster C5 driven by sample “H3” was excluded from all downstream analyses. (**C**) Dot plot indicating the top 10 marker genes per cluster (see ***Table S1***). (**D**-**F**) (**D**) Label transfer and projection based on a recently published scRNA-seq data set of P14 $T_E$, $T_M$, and $T_{EX}$ generated in acute LCMV Arm or chronic LCMV cl13 infection[2], (**E**) heatmap visualization of the top 20 cluster marker genes after label transfer, and (**F**) cluster concordance between the two scRNAseq-based cell annotations. (**G**-**H**) Pseudobulk analyses of scRNAseq data. (**G**) Volcano plot of differential expression (DE) analysis between all helped and helpless P14 $T_E$. Vertical dashed line indicates 1.5-fold change; horizontal dashed lines mark $p$=0.05 and 0.01, respectively. (**H**) Non-overlapping Venn diagram of differentially expressed genes (DEGs) by helped and helpless P14 $T_E$. (**I**) Total numbers and cellular cluster compositions for individual helped and helpless donors. (**J**-**L**) Lower resolution Leiden cluster reanalysis. (**J**) UMAP projection with lower resolution Leiden clustering combining original clusters C0, C1, and C2 (see **A**) into a single effector cluster C0* while preserving the cytotoxic C3 and memory precursor C4 clusters, (**K**) heatmap of top 40 marker genes, and (**L**) relative abundance of helped and helpless P14 $T_E$/cluster after lower-resolution re-clustering. *$p$<0.05 by unpaired Student’s $t$ test in (**L**).

#### Figure S4 (related to Figure 4)

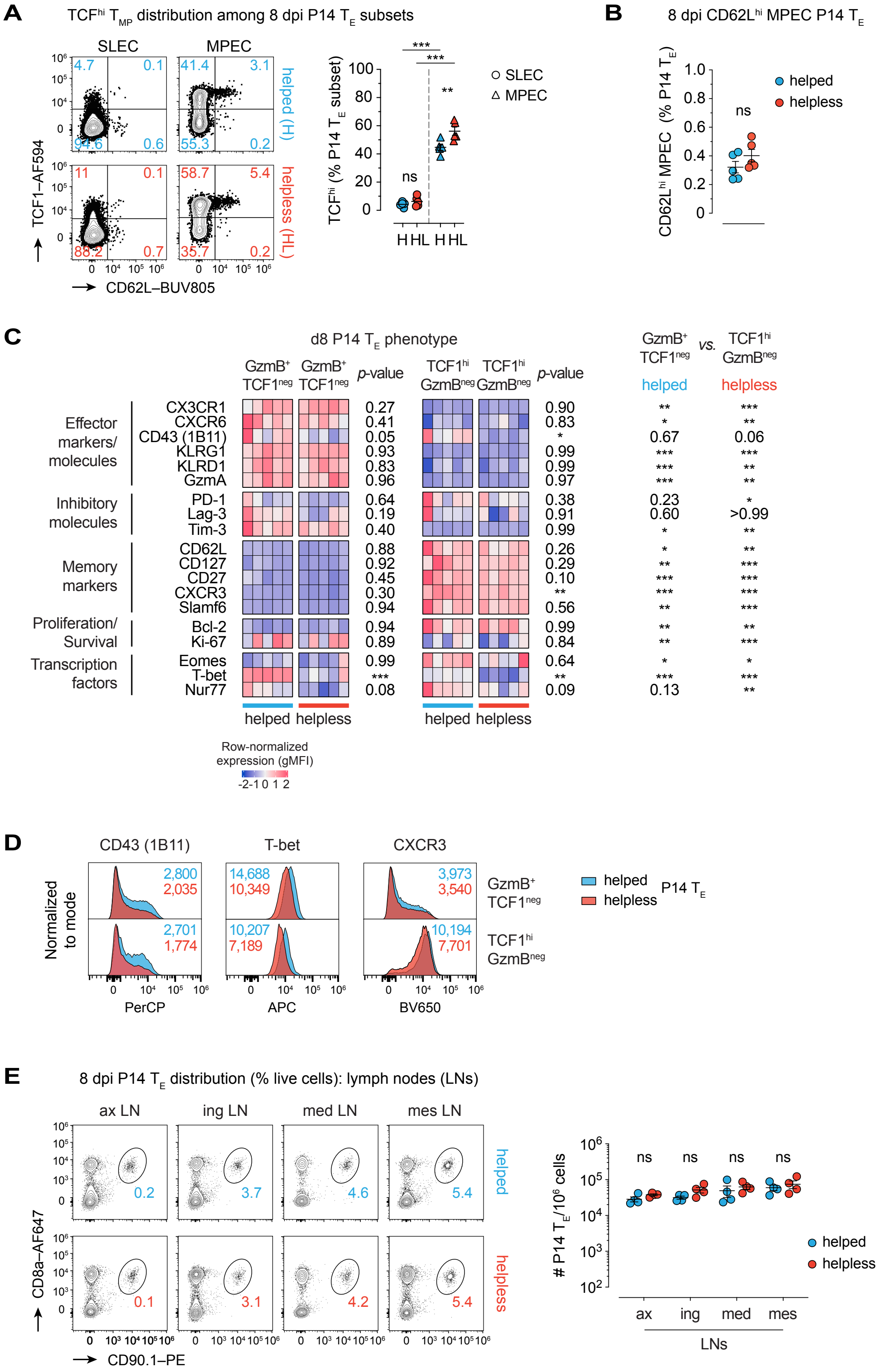

***Figure S4*** **(related to Figure 4). Minor differences between helped and helpless P14 $T_E$ at the protein and functional level**

(**A**-**B**) (Left) representative flow cytometry plots and (right) quantification of (**A**) TCF1 and (**B**) CD62L expression by splenic P14 SLEC (gated on $CD127^-KLRG1^+$ P14) and MPEC (gated on $CD127^+KLRG^-$ P14; see ***Figure 4C***) subsets. (**C**) Heatmap visualization and statistics of *Z*-score normalized row gMFI values comparing helped *vs*. helpless $GzmB^+TCF1^{neg}$ (left) or helped *vs*. helpless $TCF1^{hi}GzmB^{neg}$ (middle) splenic P14 $T_E$, as well as $GzmB^+TCF1^{neg}$ *vs*. $TCF1^{hi}GzmB^{neg}$ P14 $T_E$ (right) populations in helped or helpless mice. (**D**) Representative histogram overlays showing CD43 (clone 1B11), T-bet, and CXCR3 expression by $GzmB^+TCF1^{neg}$ and $TCF1^{hi}GzmB^{neg}$ splenic P14 $T_E$ subsets. (**E**) (Left) representative flow cytometry plots and (right) enumeration of P14 $T_E$ distribution across various LNs. Data are representative of one out of two to three independent experiments conducted at 8 dpi with n=5 mice/group/experiment in (**A**) and (**B**), and n=4 mice/group/experiment in (**C**) and (**E**). Graphs plot individual helped (blue) and helpless (red) mice and mean±SEM. *$p$<0.05, **$p$<0.01, and ***$p$<0.001 by one-way or repeated-measures ANOVA with Šídák's multiple comparisons test in (**A**) and (**C**), unpaired Student's *t* test in (**B**) and (**E**), and Mann Whitney U test in (**H**). ns, non-significant.

### Figure S5 (related to Figure 5)

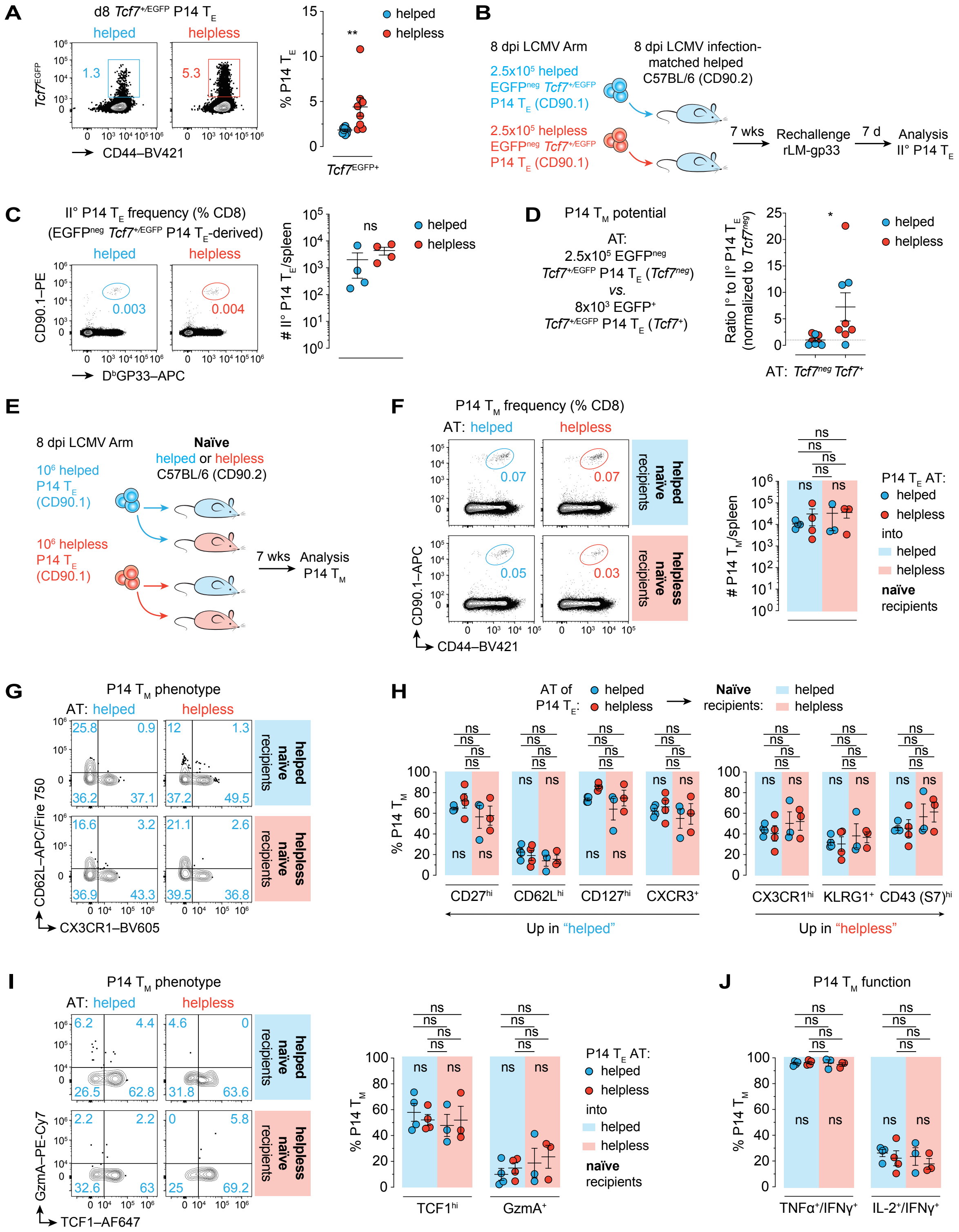

***Figure S5*** **(related to Figure 5). Helpless CD8$^+$ $T_E$ possess intrinsic $T_M$ potential that is contingent on the post-effector phase environment**

(**A**-**D**) Helped and helpless EGFP$^+$/TCF1$^+$ or EGFP$^{neg}$/TCF1$^{neg}$ P14 $T_E$ were generated by transferring naïve *Tcf7*$^{+/EGFP}$ P14 into congenic isotype-treated or transiently CD4$^+$ T cell-depleted B6 hosts followed by acute LCMV Arm infection (see ***Figures 5G*** and ***5H***). (**A**) (Left) representative flow cytometry plots and (right) quantification of *TCF7*$^{EGFP+}$ P14 $T_E$ at 8 dpi. (**B**) 2.5x10$^5$ sort-purified 8 dpi helped or helpless EGFP$^{neg}$ *Tcf7*$^{+/EGFP}$ P14 $T_E$ were transferred into congenic infection-matched helped secondary recipients and P14 $T_M$ generation as well as secondary reactivity were assessed 7 weeks later after heterologous high-dose rLM-gp33 rechallenge. (**C**) (Left) representative flow cytometry plots of splenic II° P14 $T_E$ expansion in response to rLM-gp33 and (right) enumeration. (**D**) For a comparative assessment of differential P14 $T_M$ potential by EGFP$^+$ *vs*. EGFP$^{neg}$ P14 $T_E$, the respective magnitudes of II° P14 $T_E$ expansions were divided by the corresponding input numbers of adoptively transferred I° P14 $T_E$ and normalized to average obtained ratios for EGFP$^{neg}$ adoptively transferred P14 $T_E$ (dashed line). (**E**-**J**) Enriched congenic helped or helpless 8 dpi P14 $T_E$ were adoptively transferred into naïve helped or helpless secondary recipients and splenic P14 $T_M$ maturation was assessed seven weeks later. (**E**) Experimental outline of P14 $T_E$ "criss-cross" adoptive transfers and subsequent P14 $T_M$ maturation under four distinct conditions in naïve secondary hosts (transient CD4$^+$ T cell depletion model). (**F**) (Left) representative flow cytometry plots of frequencies and (right) enumeration of adoptively transferred P14 populations derived from helped (blue circles) or helpless (red circles) P14 $T_E$ donors after maturation in respective helped (light blue background) or helpless (light red background) naïve secondary recipients. (**G**-**J**) Depicted are representative flow cytometry plots and summary analyses of (**G**-**H**) P14 $T_M$ phenotypes, (**I**) transcription factor and effector molecule expression, and (**J**) functional properties after gp33-peptide restimulation *in vitro*. Data are representative of one out of one to two independent experiments with n=8-10 mice/group/experiment in (**A**), with n=4 mice/group/experiment in (**C**), and n=3-4 mice/group in (**E**)-(**J**). Data in (**D**) are pooled helped and helpless mice from experiments outlined in ***Figures*** **5G**-**5H** and **S5B**-**S5C**. Graphs plot individual helped (blue) and helpless (red) mice and mean±SEM. *$p$<0.05, **$p$<0.01 by Mann Whitney U test in (**A**) and (**B**), by unpaired Student's $t$ test in (**C**), and by one-way ANOVA with Tukey's multiple comparisons test in (**F**), (**H**), (**I**), and (**J**). ns, non-significant.

#### Figure S6 (related to Figure 6)

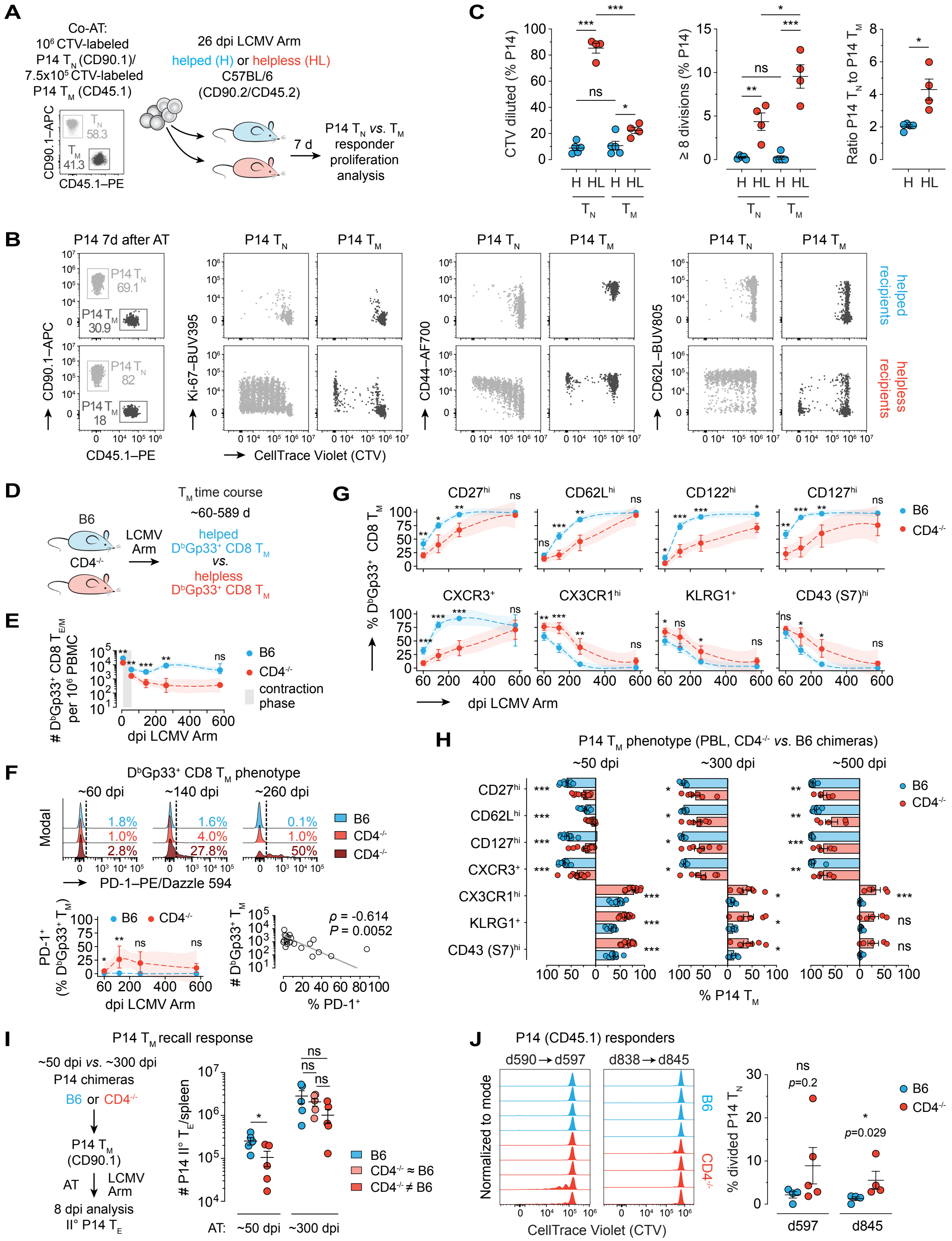

***Figure S6*** **(related to Figure 6). Prolonged cognate antigen presentation in transiently and permanently $CD4^+$ T cell-deficient mice**

(**A**-**C**) CellTrace Violet (CTV)-labeled P14 $CD90.1^+$ $T_N$ and P14 $CD45.1^+$ $T_M$ were co-transferred into congenic LCMV-immune helped or helpless B6 recipients at a ratio of 1.4:1 on d26 post LCMV Arm, recovered from spleens 7 days later, and P14 proliferation was quantified by CTV dilution. (**A**) Experimental setup for co-transfer P14 responder assay (transient $CD4^+$ T cell depletion model). (**B**) Representative flow cytometry plots of recovered helped (top row) and helpless (bottom row) splenic P14 and expression of Ki-67, CD44, and CD62L as a function of CTV-dilution. (**C**) (Left) Quantification of diluted CTV fraction (*i.e.*, percent of P14 cells that divided at least once) by P14 $T_N$ or $T_M$ responders in helped and helpless recipients, (middle) percentage of P14 $T_N$ or $T_M$ responders with ≥8 divisions in helped and helpless recipients, and (right) altered ratio of P14 $T_N$ to P14 $T_M$ 7d post transfer into helped and helpless recipients. (**D**-**J**) Assessment of $CD8^+$ $T_M$ maturation kinetics and prolonged cognate antigen presentation under conditions of permanent $CD4^+$ T cell deficiency. (**D**-**G**) (**D**) B6 (helped) or permanently $CD4^+$ T cell-deficient mice ($CD4^{-/-}$; helpless) were infected with LCMV Arm and (**E**) LCMV-specific $D^bGP_{33}^+$ $CD8^+$ $T_M$ PBMC relative numbers were followed over time from ~60-600 dpi. (**F**) (Top) representative histogram overlays of individual B6 and $CD4^{-/-}$-derived splenic $D^bGP_{33}^+$ $CD8^+$ $T_M$ featuring gradual re-expression of the activation marker PD-1 by some helpless $CD8^+$ $T_M$ over time, (bottom left) statistical quantification, (bottom right) inverse correlation of $D^bGP_{33}^+$ $CD8^+$ $T_M$ and PD-1 re-expression in helpless $CD4^{-/-}$ mice across all sampled time points post LCMV. (**G**) Kinetics of LCMV-specific $D^bGP_{33}^+$ $CD8^+$ $T_M$ phenotypes. (**H**-**I**) Helped and permanently helpless P14 $CD90.1^+$ chimeras were generated by transferring $5x10^3$ P14 $T_N$ into B6 or $CD4^{-/-}$ recipients, followed by LCMV Arm infection. (**H**) Summary of PBL P14 $T_M$ phenotypes in B6 *vs*. $CD4^{-/-}$ mice at ~50, 300, and 500 dpi featuring helpless late-phase P14 $T_M$ with either mature or more pronounced immature phenotypic profiles as indicated by scatter range. (**I**) Comparison of II° P14 $T_E$ responses by helped (B6; blue) *vs*. helpless ($CD4^{-/-}$; red) P14 $T_M$ at ~50 dpi and by helped (B6; blue) *vs*. mature-like-phenotype helpless ($CD4^{-/-}$ ≈ B6; light red) *vs*. immature-phenotype helpless ($CD4^{-/-}$ ≠ B6; red) at ~300 dpi. (**J**) (Left) representative histograms of CTV dilution and (right) summary data of percent divided by P14 CD45.1 $T_N$ responders after transfer into congenic LCMV-immune B6 or $CD4^{-/-}$ recipients at d597 or d845 post infection indicating residual cognate antigen presentation in individual permanently helpless mice (see ***Figures 6A-6C***). Data are representative of one to two independent experiments with n=4-5 mice/group in (**A**)-(**C**), n=2-8 mice/group/time point in (**D**)-(**F**), n=5-14 mice/group/time point in (**H**), n=5 mice/group/time point in (**I**), and n=4-5 mice/group/time point in (**J**). Correlation data with non-linear fit semi-log line in

(**F**) indicate Spearman's rank correlation coefficient ρ as well as *P* value and are pooled helpless mice from all memory time points sampled in (**F**)-(**G**). Graphs plot individual helped (blue) and helpless (red) mice and mean±SEM in (**C**) and (**H**)-(**J**) or mean±bootstrap 95% confidence intervals in (**E**)-(**G**). Connecting dashed lines and associated ribbons in (**E**)-(**G**) represent LOESS smoothing trends with 95% confidence intervals as calculated by the geom_smooth() function in R. *$p<0.05$, **$p<0.01$, and ***$p<0.001$ by one-way ANOVA with Šídák's multiple comparisons/mixed-effects analysis and Mann Whitney U in (**C**), unpaired Student's *t* or Mann Whitney U test in (**E**)-(**H**) and (**J**), by Spearman correlation in (**F**), and by Student's t test or one-way ANOVA with Tukey's multiple comparisons in (**I**). ns, non-significant.

#### Figure S7 (related to Figure 6)

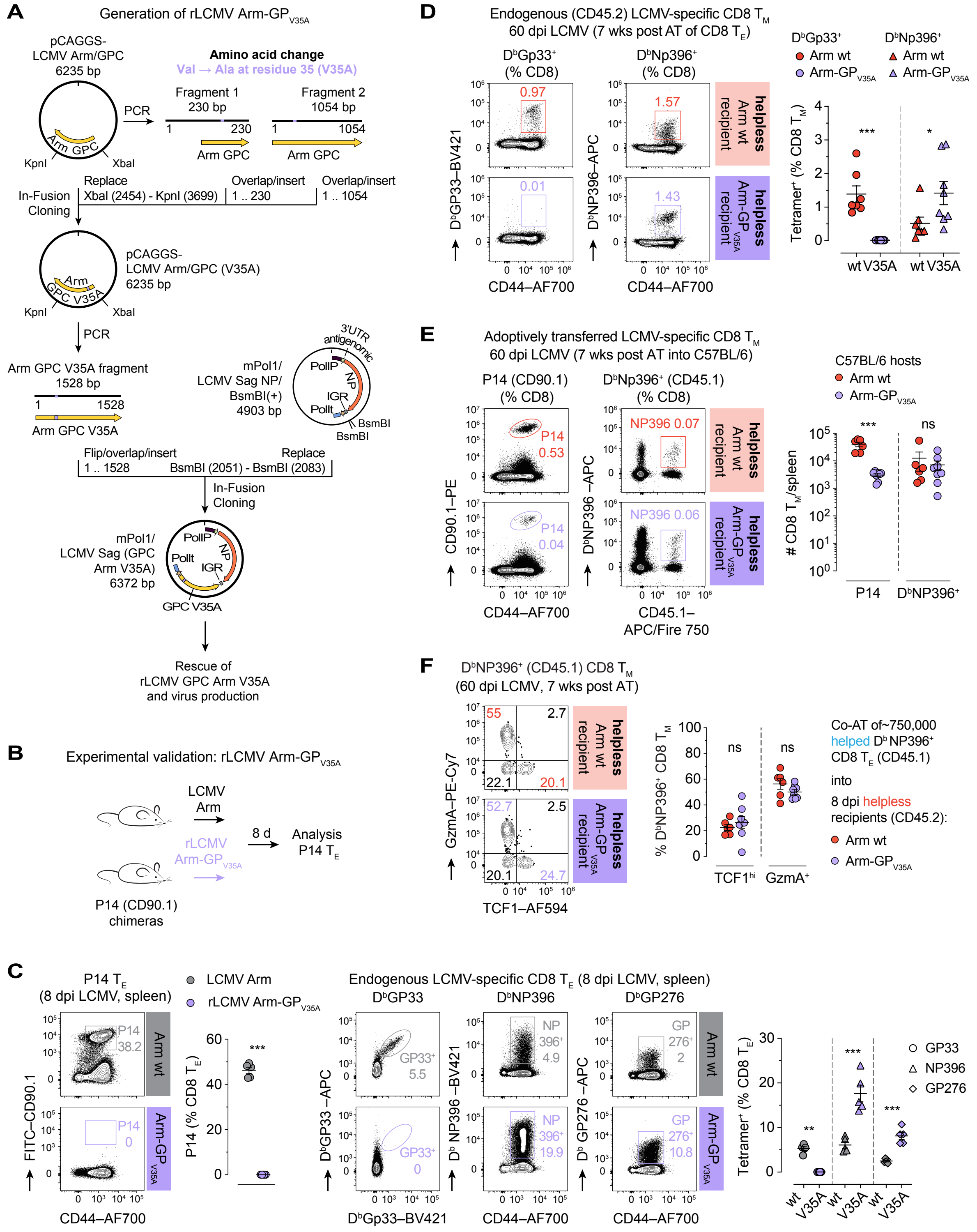

***Figure S7*** **(related to Figure 6)*.* Infection with rLCMV-GP$_{V35A}$ indicates cognate antigen presentation is causative of helpless memory**

(**A**-**C**) Generation and experimental validation of mutant rLCMV-GP$_{V35A}$ evading recognition by P14 or endogenous $D^bGp_{33}^+$ $CD8^+$ T cells. (**A**) Summary cartoon outlining generation of rLCMV-GP$_{V35A}$ by an established plasmid-based viral rescue strategy (GPC = glycoprotein precursor). (**B**) Setup for experimental validation of rLCMV-GP$_{V35A}$ *vs*. LCMV Arm following infection of P14 chimeras. (**C**) Characterization by representative flow cytometry plots and summary analyses of (left) P14 $T_E$ and (right) endogenous LCMV-specific $D^bGp_{33}^+$, $D^bNP_{396}^+$, and $D^bGP_{276}^+$ $CD8^+$ $T_E$ responses at 8 dpi with LCMV Arm *vs*. rLCMV-GP$_{V35A}$ indicating absence of recognition by P14 and $D^bGP_{33\text{-}41}$ $CD8^+$ but not $D^bNP_{396}^+$ or $GP_{276}^+$ $CD8^+$ T cells. (**D**-**F**) 8 dpi helped splenic P14 $CD90.1^+$ $T_E$ were generated in $CD45.1^+$ B6 mice and transferred into 8 dpi congenic ($CD90.2^+$/$CD45.2^+$) helpless secondary hosts infected with LCMV Arm wt (red) or with rLCMV-GP$_{V35A}$ (purple) as outlined in ***Figure 6F***. (**D**) (Left) representative flow cytometry plots and (right) quantification of splenic $D^bGp_{33}^+$ and $D^bNP_{396}^+$ $CD45.2^+$ $CD8^+$ $T_M$ in LCMV Arm wt- or LCMV Arm-GP33$_{V35A}$-immune B6 hosts 7 weeks after P14 $T_E$ adoptive transfers demonstrating lack of endogenous $D^bGp_{33}^+$ $CD8^+$ $T_M$ and slightly increased percentages of $D^bNP_{396}^+$ $CD8^+$ $T_M$ in rLCMV Arm-GP33$_{V35A}$-immune mice. (**E**) (Left) representative flow cytometry plots and (right) enumeration of splenic P14 $CD90.1^+$ and $D^bNP_{396}^+$ $CD45.1^+$ $CD8^+$ $T_M$ derived from adoptively co-transferred P14 or $D^bNP_{396}^+$ $CD8^+$ $T_E$, respectively, showing moderately increased P14 but equivalent $D^bNP_{396}^+$ $CD8^+$ $T_M$ in helpless rLCMV-GP$_{V35A}$- *vs*. LCMV Arm wt-immune hosts. (**F**) Shown are (left) representative flow cytometry plots and (right) summary of $T_M$ phenotypes by $D^bNP_{396}^+$ $CD45.1^+$ $CD8^+$ $T_M$ derived from $D^bNP_{396}^+$ $CD45.1^+$ $CD8^+$ $T_E$ adoptively co-transferred together with P14 $T_E$ into congenic 8 dpi helpless LCMV Arm- or rLCMV-GP$_{V35A}$-immune B6 recipients. Data are representative of one out of at least two independent experiments with n=5 mice/group/experiment in (**C**) and n=5-8 mice/group/experiment in (**D**-**F**). Graphs plot individual mice with respective group color association as indicated and mean±SEM. *$p$<0.05, **$p$<0.01, and ***$p$<0.001 by unpaired Student's $t$ or Mann Whitney U test. ns, non-significant.

**Supplemental references**


1. Milner, J.J., Nguyen, H., Omilusik, K., Reina-Campos, M., Tsai, M., Toma, C., Delpoux, A., Boland, B.S., Hedrick, S.M., Chang, J.T., and Goldrath, A.W. (2020). Delineation of a molecularly distinct terminally differentiated memory CD8 T cell population. Proc Natl Acad Sci U S A *117*, 25667-25678. 10.1073/pnas.2008571117.
2. Giles, J.R., Ngiow, S.F., Manne, S., Baxter, A.E., Khan, O., Wang, P., Staupe, R., Abdel-Hakeem, M.S., Huang, H., Mathew, D., et al. (2022). Shared and distinct biological circuits in effector, memory and exhausted CD8(+) T cells revealed by temporal single-cell transcriptomics and epigenetics. Nat Immunol *23*, 1600-1613. 10.1038/s41590-022-01338-4.
